# Caspase-8, RIPK1, and RIPK3 Coordinately Regulate Retinoic Acid-Induced Cell Differentiation and Necroptosis

**DOI:** 10.1101/156901

**Authors:** Masataka Someda, Shunsuke Kuroki, Makoto Tachibana, Shin Yonehara

**Affiliations:** Graduate School of Biostudies, Kyoto University, Kyoto 606-8501, Japan.; The Institute for Enzyme Research, The University of Tokushima, Tokushima 770-8503, Japan.

## Abstract

Caspase-8, which is essential for death receptor-mediated apoptosis, inhibits necroptosis by suppressing the function of RIPK1 and RIPK3 to activate MLKL. We show that knockdown of *caspase-8* expression in embryoid bodies derived from ES cells markedly enhances retinoic acid (RA)-induced cell differentiation and necroptosis, both of which are dependent on *Ripk1* and *Ripk3.* RA treatment obviously enhanced the expression of RA-specific genes having a retinoic acid response element *(RARE)* to induce cell differentiation, and induced marked expression of RIPK1, RIPK3 and MLKL to stimulate necroptosis. *Caspase-8* knockdown induced RA receptor (RAR) to form a complex with RIPK1 and RIPK3 in the nucleus, and RAR interacting with RIPK1 and RIPK3 showed much stronger binding activity to *RARE* than RAR without RIPK1 or RIPK3. In Caspase-8-deficient mouse embryos, expression of RA-specific genes was obviously enhanced. Thus, caspase-8, RIPK1, and RIPK3 regulate RA-induced cell differentiation and necroptosis both *in vitro* and *in vivo.*

## Introduction

Apoptosis is an important physiological cell suicide mechanism essential for normal embryonic development and the maintenance of homeostasis in adult tissues. Caspases, members of the cysteine protease family, play an essential role in the induction of apoptosis (Alnemri, 1997; Chinnaiyan et al., 1997). Two major pathways of mammalian apoptosis are known, a pathway through mitochondria (intrinsic pathway) and a pathway through death receptors (extrinsic pathway) (Green, 2005). Both pathways require the activation of a variety of caspases, which are known to be pro-enzymes activated by cleavage at aspartate residues upon death signaling (Salvesen and Dixit, 1997).

Caspase-8 was originally identified as an initiator caspase and mainly functions in the death receptor pathway of apoptosis. Upon ligation of a death receptor such as Fas ( Yonehara et al., 1989; Itoh et al., 1991), caspase-8 is recruited to a complex known as a death-inducing signaling complex (DISC) together with other factors including Fas, Fas-associated death domain (FADD) (Muzio et al., 1996). Within the complex, proximity-induced auto-cleavage through homo-oligomerization/dimerization catalytically activates caspase-8. The activated caspase-8 transmits the death signal mainly to executor caspases including caspase-3, which then cleave various cellular proteins to complete the apoptosis-inducing process (Thornberry and Lazebnik, 1998). Caspase-8 is unique, with associated critical activities not only to induce apoptosis but also to suppress death receptor-mediated necroptosis ( Varfolomeev et al., 1998; Oberst et al., 2011). Caspase-8 inhibits necroptosis by suppressing the function of receptor interacting protein kinase 1 (RIPK1 or RIP1) and RIPK3 (or RIP3) (Hitomi et al., 2008; He et al., 2009; Zhang et al., 2009) to activate MLKL, an executer molecule of necroptosis (Wang et al., 2014). Disruption of *caspase-8 (Casp8)* expression causes embryonic lethality in mice around embryonic day 11.5 (E11.5) (Varfolomeev et al., 1998; Sakamaki et al., 2002), which is rescued by depletion of *Ripk3* or *mlkl,* indicating that the embryonic lethality is caused by activation of necroptosis (Kaiser et al., 2011; Alvarez-Diaz et al., 2016).

Retinoic acid (RA), which is a metabolic product of vitamin A, is a well-established signaling molecule that plays essential roles in vertebrate embryonic body shaping and organogenesis, tissue homeostasis, cell proliferation, differentiation, and apoptosis by regulating the expression of RA-specific target genes (Mark, 2005; Duester, 2008; Rhinn and Dolle, 2012). RA binds to a transcription complex in nucleus, which includes a pair of ligand-activated transcription factors composed of the RA receptor (RAR)-retinoic X receptor (RXR) heterodimer, to induce transcription of RA-specific genes. There are three RAR genes *(Rara, Rarb* and *Rarg)* and three RXR genes *(Rxra, Rxrb* and *Rxrg),* and the heterodimeric pair binds to a DNA sequence called a retinoic acid-response element (RARE) (Dolle et al., 1989, Mangelsdorf, 1990, Mangelsdorf, 1991). Genes containing RARE in their promoters are known to be involved in diverse but interrelated biological processes, such as embryogenesis, growth, and differentiation (Durand et al., 1992). Following the successful application of RA in the differentiation therapy of acute promyelocytic leukemia (APL), regulation of RA signaling was also related to differentiation, proliferation or apoptosis of tumor cells (Wang and Chen, 2008; Ablain et al., 2014).

## Results

### Knockdown of *Caspase-8* Expression Evidently Enhances RA-Induced Cell Differentiation

The roles of caspase-8 on growth, viability and differentiation were investigated in ES cells by utilizing a tetracycline/doxycycline (Dox)-inducible short hairpin RNA (shRNA) expression (Tet-On) system (Kobayashi and Yonehara, 2009) specific for *Casp8* (sh*Casp8)* (Figure 1A). While *Casp8* expression was clearly down-regulated by treatment with Dox in ES cells with the Tet-On sh*Casp8* system (Tet-On *shCasp8* ES cells) (Figure 1B), significant effects on neither growth nor viability were observed in the ES cells after treatment with Dox. Furthermore, *in vitro* neural differentiation induced by embryoid body (EB) formation (Watanabe et al., 2005) was comparable in ES cells irrespective of whether *Casp8* expression was down-regulated or not. Interestingly, however, cell differentiation was remarkably enhanced in EBs derived from *Casp8* KD ES *(Casp8* KD ES) cells after 6-day treatment with RA (Figures 1C, Figure 1 - figure supplement 1A). *Oct3/4,* a marker of undifferentiated cells, was strongly down-regulated in RA-treated EBs derived from *Casp8* KD ES cells (Figure 1D), and the expression levels of neural differentiation markers, *Nestin* and *Tuj1,* were up-regulated in RA-treated EBs derived from *Casp8* KD ES cells relative to control *shGFP* ES cells (Figure 1E). While the expression level of *Casp8* was also up-regulated in the RA-treated EBs, expression of *Casp8* was almost entirely diminished for the entire time in *Casp8* KD EBs (Figure 1 - figure supplement 1B). Thus, KD of *Casp8* expression in ES cells markedly enhances RA-induced neural differentiation.

**Figure 1.**
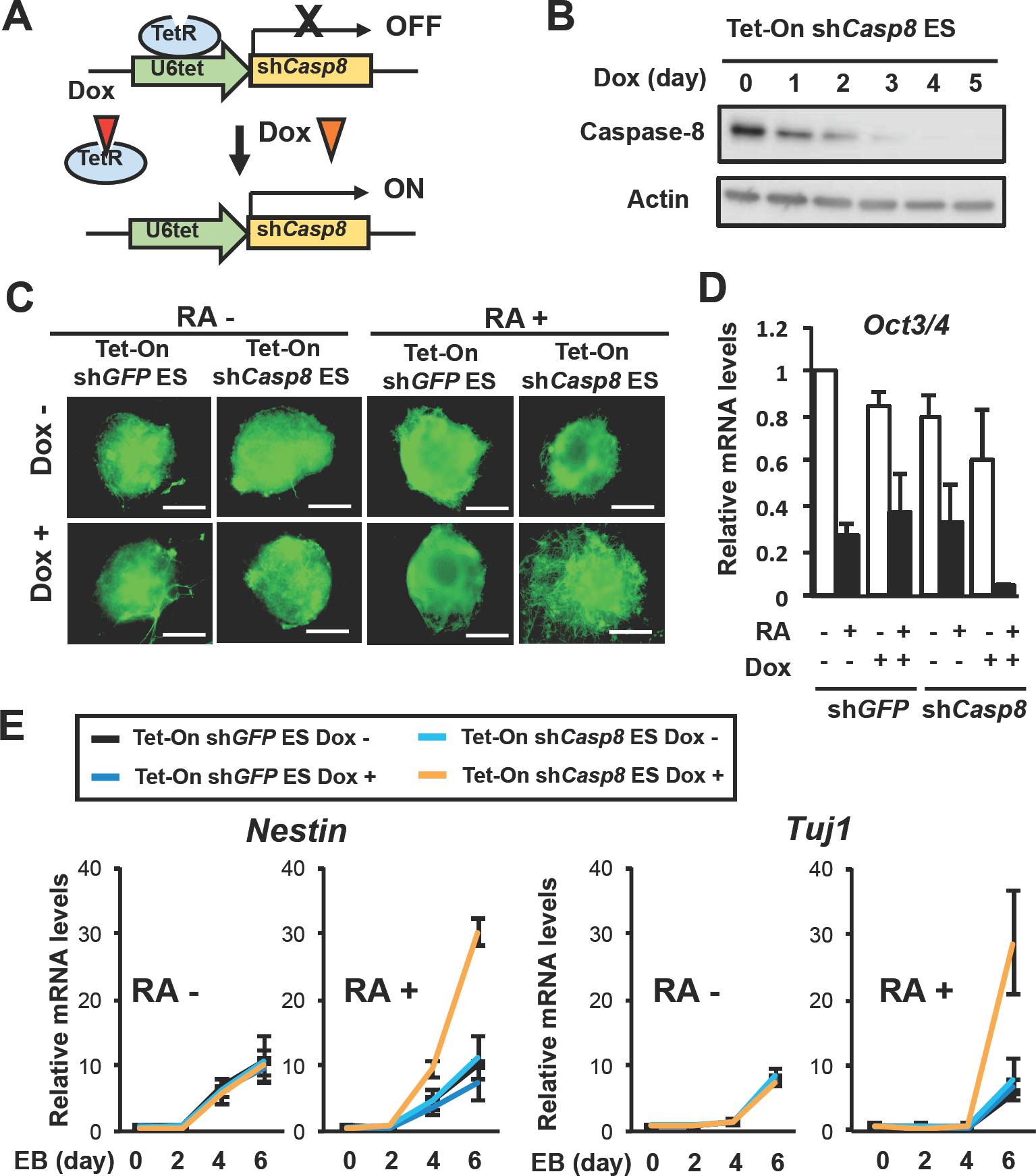
Knockdown of *Casp8* in ES cells enhances RA-induced cell differentiation. (A) Dox-inducible (Tet-On) sh*Casp8*-expression system in ES cells. TetR, tetracycline repressor; and U6tet, mouse *U6* promoter joining the tetracycline operator.
(B) Validation of induced knockdown of *Casp8* expression in Tet-On sh*Casp8* ES cells by western blot analysis after treatment with 1μg/ml Dox for indicated days.
(C) Neuronal differentiation of Dox (1 μg/ml)-treated or -untreated Tet-On sh*Casp8* and Tet-On sh*GFP* ES cells was analyzed after 6 days formation of EBs. EBs were treated with or without 1 μM RA for last 4 days. Fluorescence microscopy analysis was performed after staining with anti-Tuj 1 antibody. Scale bars, 200 μm.
(D) qRT-PCR analysis of *Oct3/4* was carried out using EBs derived from Tet-On sh*Casp8* and Tet-On sh*GFP* ES cells after 6 days formation of EBs in the presence or absence of 1 μg/ml Dox. EBs were treated with or without 1 μM RA for last 4 days. Representative data are shown as means ± SEM (n=3).
(E) qRT-PCR analysis of *Nestin* and *Tuj1* was carried out using Tet-On sh*Casp8* and Tet-On sh*GFP* ES cells at the indicated times after formation of EBs in the presence or absence of 1 μg/ml Dox. EBs were cultured with or without 1 μM RA for last 4 days. Representative data are shown as means ± SEM (n=3).

**Figure 1 – figure supplement 1.**
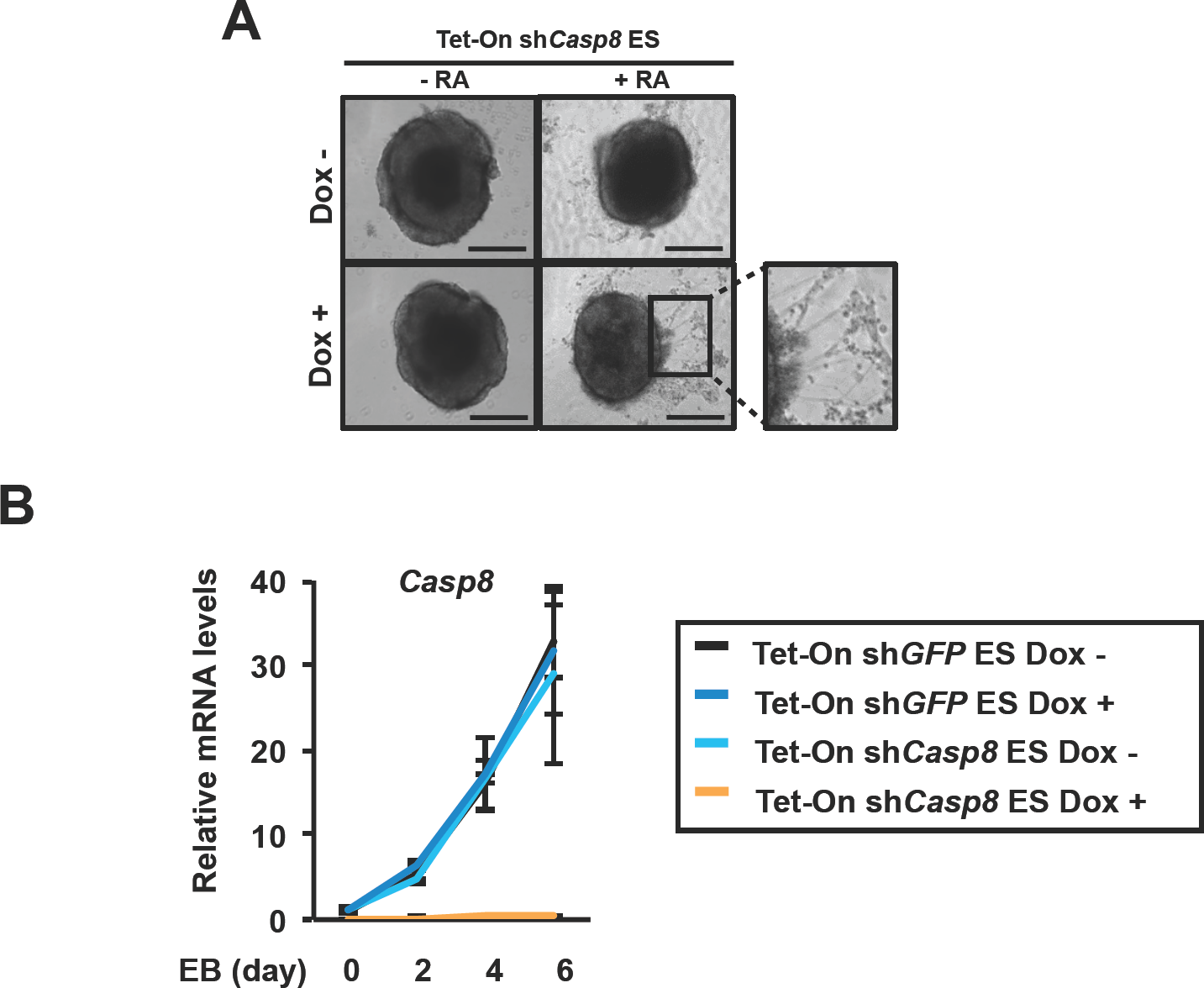
RA-induced neuronal differentiation of Tet-On sh*Casp8* ES cells. (A) Neuronal differentiation of Dox (1 μg/ml)-treated or -untreated Tet-On sh*Casp8* ES cells was analyzed under a phase-contrast microscope after 6 days formation of EBs. EBs were treated with or without 1 μM RA for last 4 days. Scale bars, 200 μm.
(B) qRT-PCR analysis of *Casp-8* in Tet-On sh*Casp8* and Tet-On shGFP ES cells at the indicated times after formation of EBs in the presence or absence of 1 μg/ml Dox. EBs were cultured with or without 1 μM RA after 2 days formation of EBs. Representative data are shown as means ± SEM (n=3).

### Knockdown of *Caspase-8* or *Fadd* Markedly Enhances RA Signaling

We then analyzed expression levels of RA-induced genes, *Crabp2, Hoxb1, Cyp26a1* and *Rarb,* which expressions were under the control of RARs and a *RARE* in the respective promoter regions of these genes (Rossant et al., 1991; Astrom et al., 1994, Ogura and Evans, 1995; Loudig et al., 2000). Quantitative real time polymerase chain reaction (qRT-PCR) and dual-luciferase reporter analyses revealed that expression levels of all the RA-specific genes and RARE-dependent transcription of luciferase were dramatically elevated in *Casp8* KD ES cells treated with 1μM RA (Figures 2A,B). The enhancement of RA-specific genes expression in *Casp8* KD ES cells was significantly induced by even 10 nM RA (Figure 2 - figure supplement 1A). Thus, caspase-8 was shown to suppress evident activation of RA signaling in ES cells. While one of the RA-induced genes, which expression were enhanced by KD of *Casp8,* was *Rarb* (RA receptor β), the expression levels of other types of RARs than *Rarb,* such as *Rara, Rarg* and *Rxra* (Zelent et al., 1989; Mattei et al., 1991), were not influenced by KD of *Casp8* (Figure 2 - figure supplement 1B). Thus, caspase-8 suppresses evident activation of RA signaling in ES cells and mechanisms other than a general increase of *RARs* expression might be involved in the enhancement of RA signaling in the absence of *Casp8* expression.

**Figure 2.**
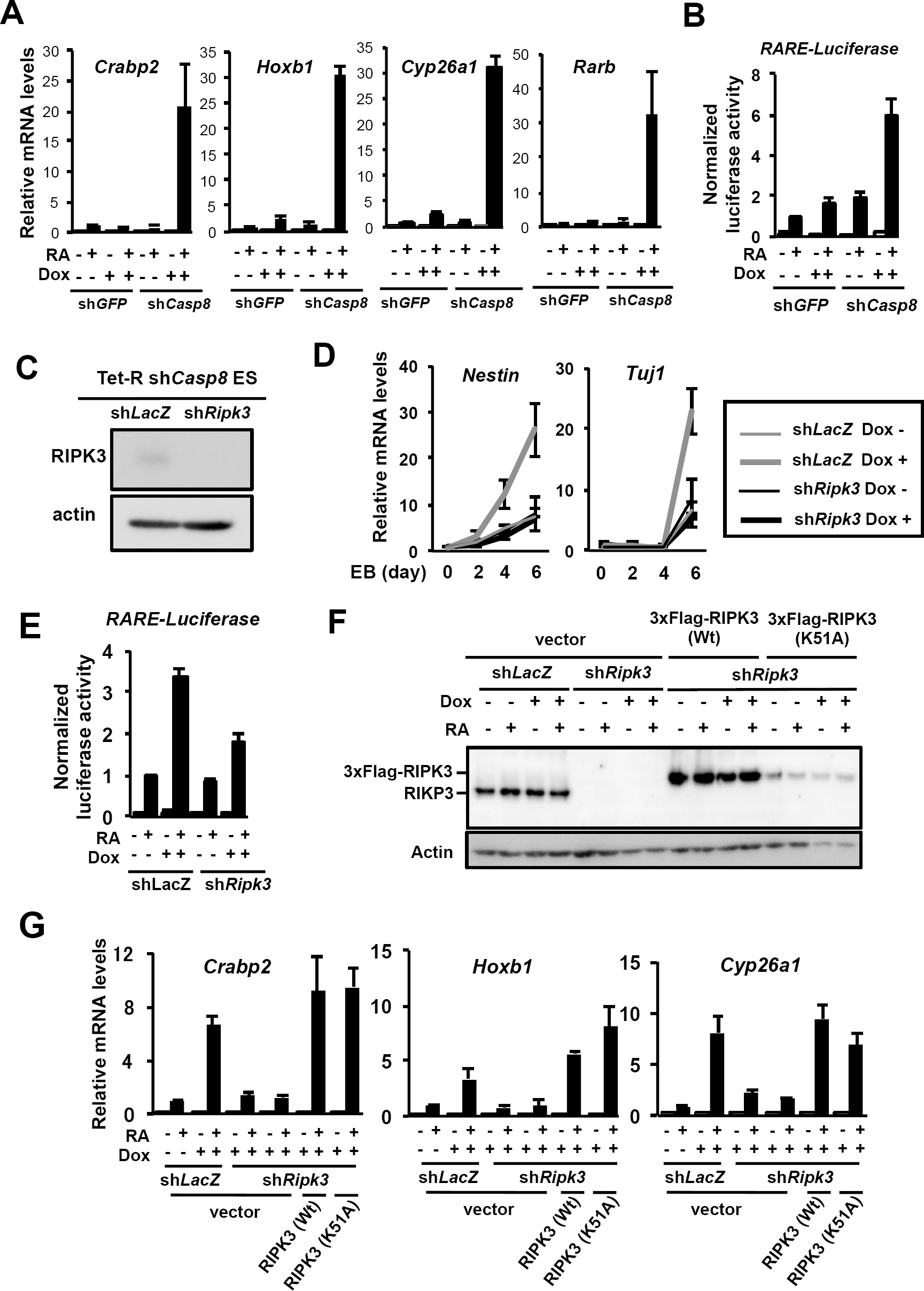
Knockdown of *Casp8* expression in ES cells markedly enhances RA signaling dependently on RIPK3. (A) Tet-On sh*Casp8* and Tet-On sh*GFP* ES cells were cultured for 4 days with or without 1 μg/ml Dox and then treated with or without 1 μM RA for 24 h in the presence or absence of Dox. Subsequently, expression levels of RA-inducible genes, *Crabp2, Hoxb1, Cyp26a1* and *Rarb,* were analyzed by qRT-PCR. Representative data are shown as means ± SEM (n=3).
(B) Dual-luciferase assay for *RARE* was performed using Tet-On sh*Casp8* or Tet-On sh*GFP* ES cells after treatment with or without 1 μM RA for 24 h in the presence or absence of 1 μg/ml Dox. Representative data are shown as means ± SEM (n=3).
(C) Western blot analysis of RIPK3 expression was carried out in Tet-On sh*Casp8* ES cells expressing sh*LacZ* or sh*Ripk3*. Actin was detected as a control.
(D) qRT-PCR analysis of *Nestin* and *Tuj1* expression was performed using EBs derived from Tet-On sh*Casp8* ES cells expressing sh*LacZ* or sh*Ripk3* after 6 days formation of EBs in the presence or absence of 1 μg/ml Dox. EBs were treated with or without 1 μM RA for last 4 days. Representative data are shown as means ± SEM (n=3).
(E) Dual-luciferase assay for *RARE* was carried out using Tet-On sh*Casp8* ES cells expressing shLacZ or sh*Ripk3* after treatment with or without 1 μM RA for 24 h in the presence or absence of 1 μg/ml Dox. Representative data are shown as means ± SEM (n=3).
(F and G) Tet-On sh*Casp8* P19 cells expressing sh*LacZ* or sh*Ripk3* were infected with lentiviral vectors carrying 3xFlag-Wt *Ripk3* or K51A *Ripk3.* These cells were cultured with or without 1 μg/ml Dox for 5 days, and then treated with or without 1 μM RA for 24 h in the presence or absence of Dox. Subsequently, western blot analysis for RIPK3 (F) and qRT-PCR analysis for RA-inducible genes (G) were carried out. vector, an empty vector. Representative data are shown as means ± SEM (n=3).

**Figure 2 - figure supplement 1.**
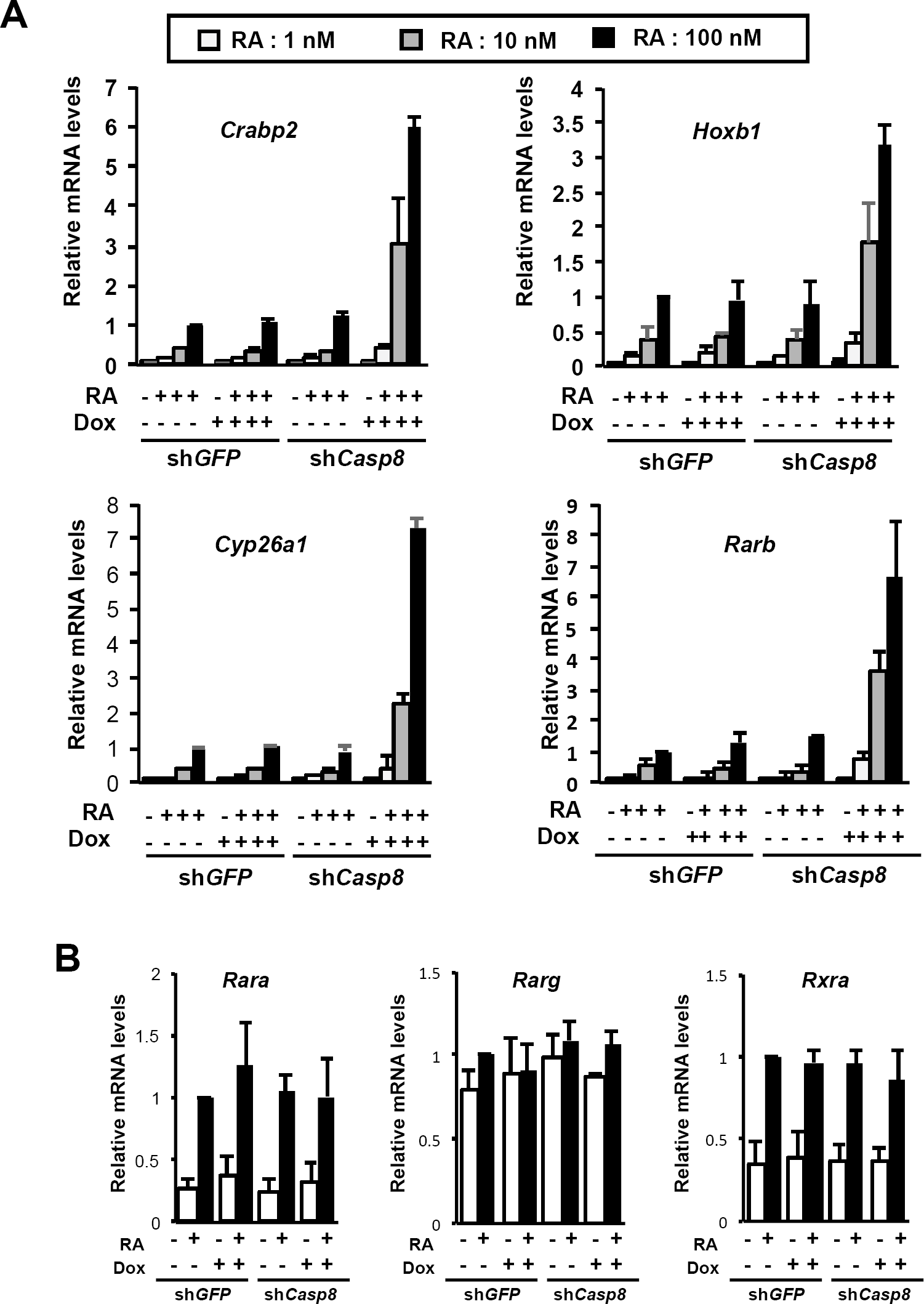
RA signaling in *Casp8* KD ES cells. (A) Tet-On sh*Casp8* and Tet-On sh*GFP* ES cells were cultured for 4 days with or without 1 μg/ml Dox and then treated with or without 1 nM, 10 n M, 100 nM or 1 μM RA for 24 h in the presence or absence of Dox. Subsequently, expression levels of RA-inducible genes, *Crabp2, Hoxbl, Cyp26al* and *Rarb,* were analyzed by qRT-PCR. Representative data are shown as means ± SEM (n=3).
(B) qRT-PCR analysis of RA receptors, *Rcirci. Rare* and *Rxra,* was carried out using Tet-On sh*Casp8* and Tet-On sh*GFP* ES cells after treatment with or without 1 μM RA for 24 h in the presence or absence of 1 μg/ml Dox. Representative data are shown as means ± SEM (n=3).

The enhancement of RA signaling as well as RA-induced cell differentiation in the absence of *Casp8* expression was observed in not only mouse ES cells but also mouse embryonic carcinoma cell line P19 (Figure 2 - figure supplement 2). In addition, KD of caspase-8 expression in human RA-sensitive cancer cell lines, SK-N-SH and HL60, clearly enhanced RA signaling and RA-induced differentiation into neural cells expressing *TUJ1* and monocytes expressing *CD11b,* respectively (Figure 2 - figure supplement 3). Thus, caspase-8 suppresses marked activation of RA signaling in not only mouse ES cells but also mouse embryonic carcinoma cells and human cancer cells.

To find out whether the protease activity and/or cleavage-associated activation of caspase-8 are necessary for inhibition of the evident activation of RA signaling, rescue experiments were carried out using exogenous expression of KD-resistant wild-type (Wt) or two kinds of *Casp8* mutants, CS and DE, which hold a Cys to Ser mutation in the protease domain (C362S) and 7 Asp to Glu mutations in expected cleavage sites of caspase-8 itself and other caspases, respectively (Kikuchi et al., 2012). While expressions of Wt and DE *Casp8* significantly inhibited the evident activation of RA signaling by *Casp8* KD, DE mutant did not really inhibit the activation of RA signaling in Tet-On *shCasp8* P19 cells (Figure 2 - figure supplement 4A,B). These results show that protease activity of procaspase-8 is indispensable but cleavage-associated activation of procaspase-8 is not necessary for inhibition of the marked activation of RA signaling. We then analyzed the KD effect of *Fadd*, a component of the death-inducing signaling complex (DISC) (Chinnaiyan et al., 1997) and the necrosome (He et al., 2009) using Tet-On sh*Fadd* ES cells, indicating that KD of *Fadd* expression induced evident activation of RA-induced neural differentiation and RA-induced genes expression (Figure 2 - figure supplement 4C,D). Taken together, the protease activity of procaspase-8 and FADD prevent marked activation of RA signaling.

**Figure 2 - figure supplement 2.**
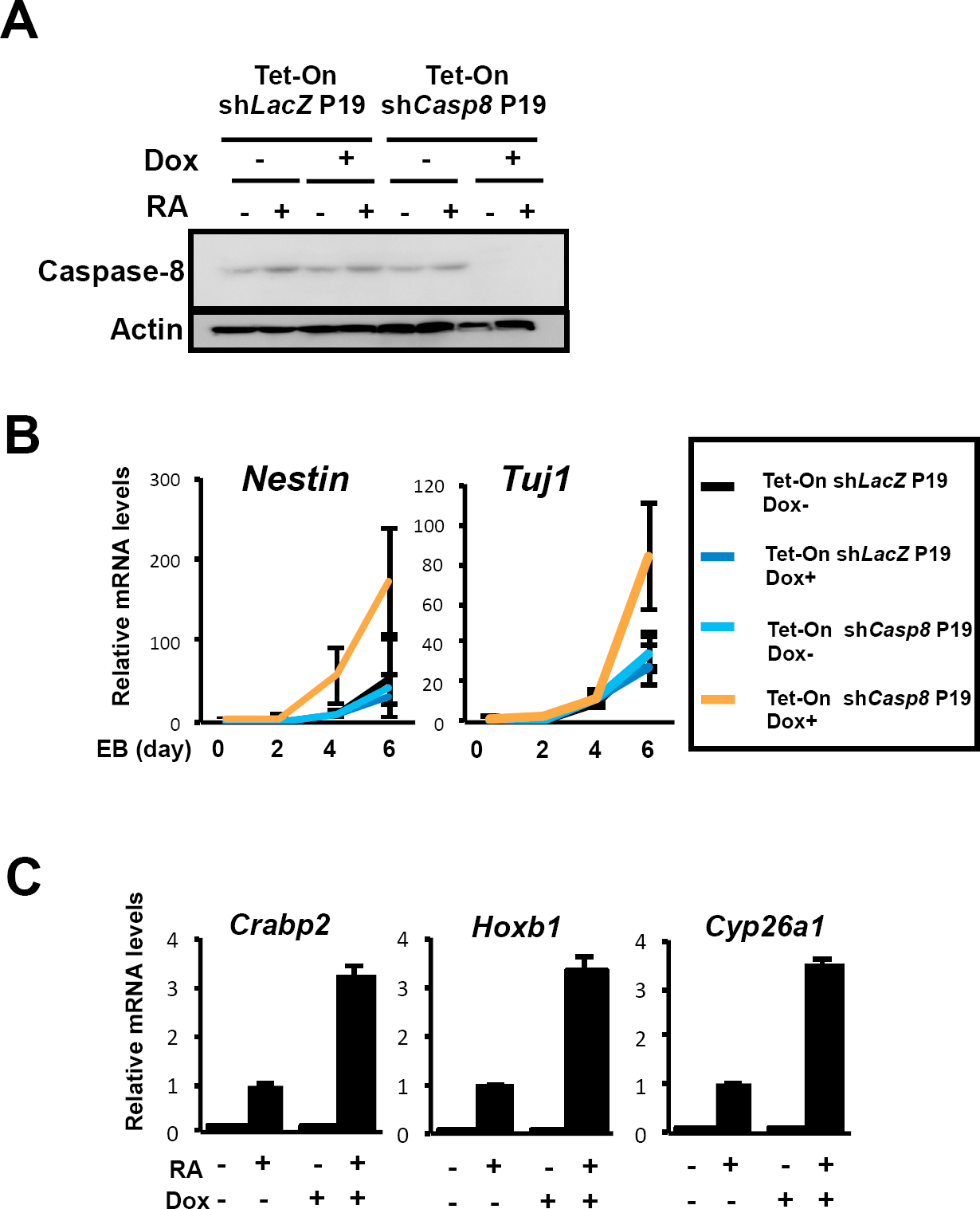
RA-induced differentiation of *Casp8* KD P19 cells. (A) Tet-On sh*Casp8* and Tet-On sh*LacZ* P19 cells were cultured with or without 1 μg/ml Dox for 4 days and then treated with 1 μM RA for 24 h in the presence or absence of Dox. Subsequently, western blot analysis with anti-caspase-8 antibody was performed. Actin was detected as a control.
(B) qRT-PCR analysis of *Nestin* and *Tuj1* was carried out using Tet-On sh*Casp8* or Tet-On sh*LacZ* P19 cells at the indicated times after formation of EBs in the presence or absence of 1 μg/ml Dox. EBs were cultured with or without 1 μM RA after 2 days formation of EBs. Representative data are shown as means ± SEM (n=3).
(C) qRT-PCR analysis of RA-inducible genes, *Crabp2, Hoxbl* and *Cyp26al*, was carried out using Tet-On sh*Casp8* P19 cells after treatment with or without 1 μM RA for 24 h in the presence or absence of 1 μg/ml Dox. Representative data are shown as means ± SEM (n=3).

**Figure 2 - figure supplement 3.**
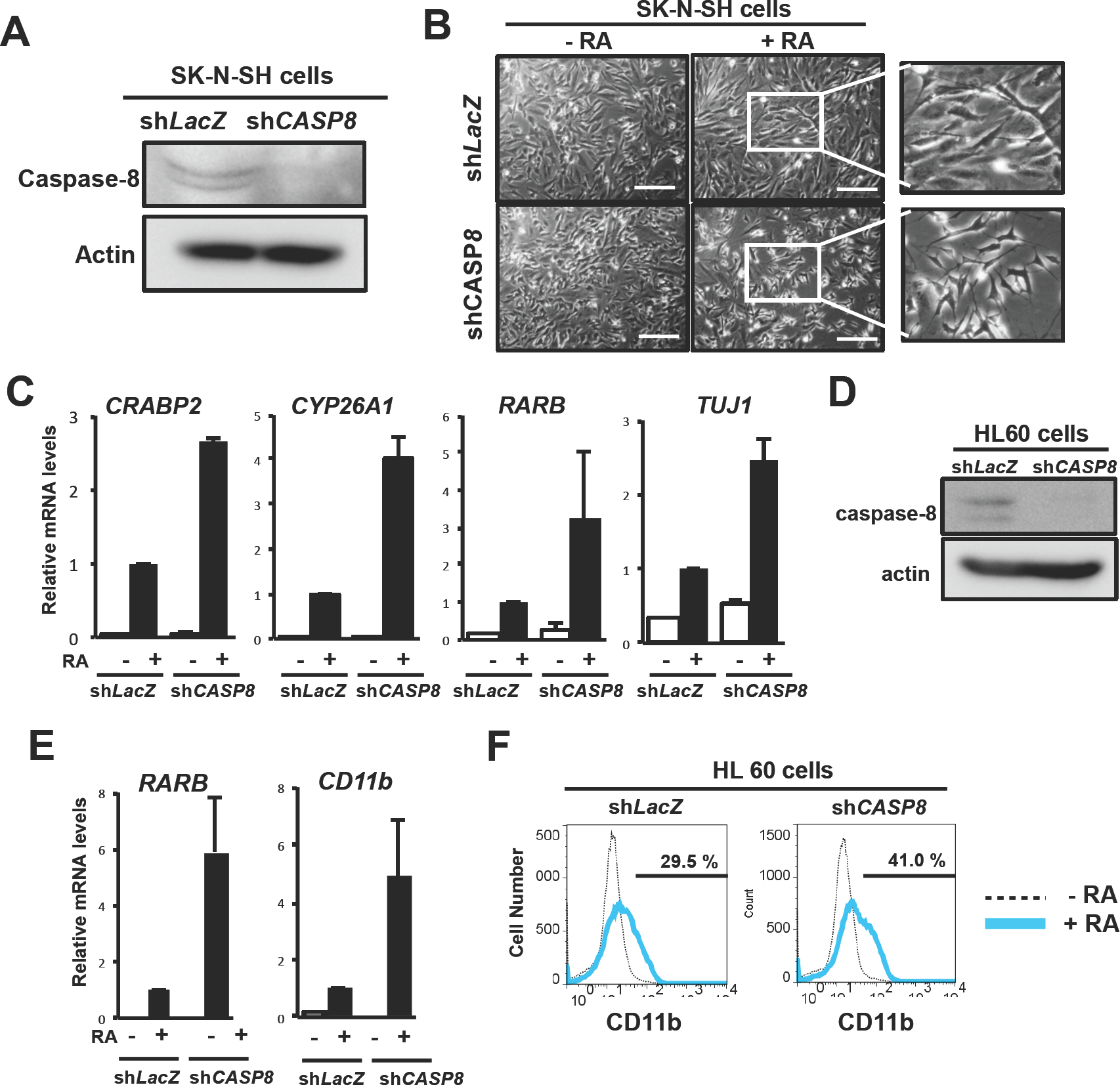
RA-induced differentiation of *Casp8* KD SK-N-SH and HL60 cells2. (A) Western blot analysis with anti-caspase-8 antibody was carried out using human neuroblastoma-derived SK-N-SH cells expressing sh*CASPS* or sh*LacZ.* Actin was detected as a control.
(B) Neuronal differentiation of SK-N-SH cells expressing sh*CASP8* or sh*LacZ* was induced by treatment with 1 μM RA for 3 days, and observed by phase-contrast microscopy. Scale bar, 200 μm.
(C) qRT-PCR analysis of *TUJ1* and RA-induced genes, *CRABP2, CYP26A1,* and *RARB,* was carried out using SK-N-SH cells expressing sh*LacZ* or sh*CASP8* after treatment with or without 1 μM RA for 24 h. Representative data are shown as means ± SEM (n=3).
(D) Human promyelocytic leukemia-derived HL60 cells expressing sh*LacZ* or sh*CASP8* were monitored for caspase-8 depletion by western blot analysis with an anti-caspase-8 antibody. Actin was detected as a control.
(E) qRT-PCR analysis of *Rarb* and *CDllb* expression was carried out using HL60 cells expressing sh*LacZ* or sh*CASP8* before and after treatment with or without 1 μM RA for 3 days. Representative data are shown as means ± SEM (n=3).
(F) HL60 cells expressing sh*LacZ* or sh*CASP8* were treated with or without 1 μM RA for 3 days, stained with an FITC-conjugated anti-CD11b antibody and subjected to flow cytometric analysis.

**Figure 2 - figure supplement 4.**
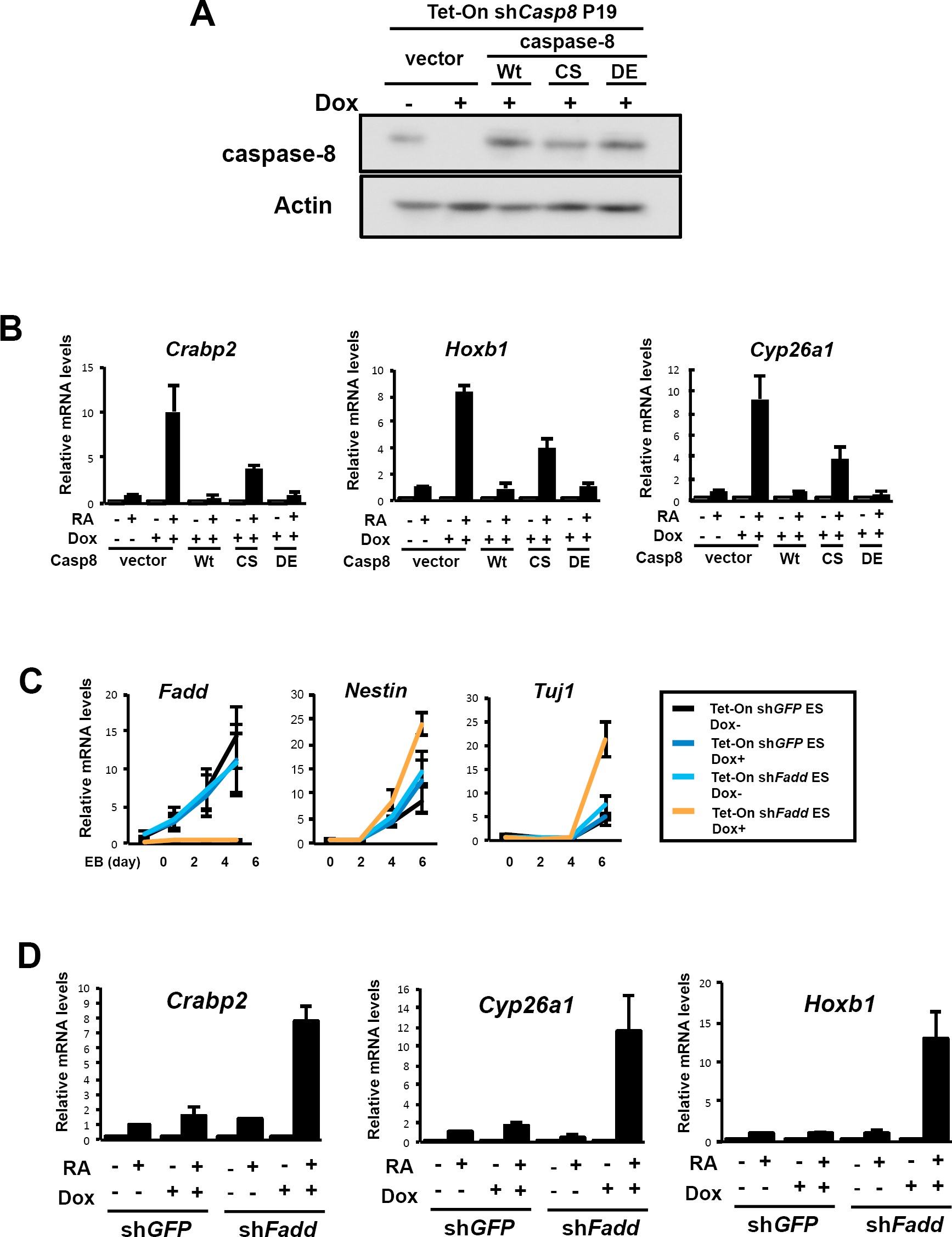
Protease activity of procaspase-8 and FADD regulate the evident activation of RA signaling. (A) Western blot analysis with an anti-caspase-8 antibody was carried out using Tet-On sh*Casp8* P19 cells expressing the following *Casp8* cDNA constructs: Wt; a protease-inactive mutant (CS) with a C362S mutation in the protease domain; or a mutant lacking putative cleavage sites (DE) with 7 Asp to Glu mutations in expected cleavage sites of caspase-8 itself and other caspases (Kikuchi et al. 2012). All the *Casp8* cDNA constructs carried silent mutations in the sh*Casp8* target sequence. After treatment with 1 mg/ml Dox for 4 days, western blot analysis was performed. Actin was detected as a control.
(B) Tet-On sh*Casp8* p19 cells were cultured for 4 days with or without 1 μg/ml Dox and then treated with or without 1 μM RA for 24 h in the presence or absence of Dox. Subsequently, expression levels of RA-inducible genes, *Crabp2, Hoxb1,* and *Cyp26a1,* were analyzed by qRT-PCR. vector, an empty vector. Representative data are shown as means ± SEM (n=3).
(C) qRT-PCR analysis of *Fadd, Nestin* and *Tuj1* was carried out in Tet-On sh*GFP* and Tet-On sh*Fadd* ES cells at the indicated times after formation of EBs in the presence or absence of 1 μg/ml Dox. EBs were cultured with or without 1 μM RA after 2 days formation of EBs. Representative data are shown as means ± SEM (n=3).
(D) Tet-On sh*Fadd* and Tet-On sh*GFP* ES cells were cultured for 4 days with or without 1 μg/ml Dox and then treated with or without 1 μM RA for 24 h in the presence or absence of Dox. Subsequently, qRT-PCR analysis of RA-induced genes, *Crabp2, Hoxb1* and *Cyp26a1,* was performed. Representative data are shown as means ± SEM (n=3).

### RIPK1 and RIPK3 but MLKL Are Involved in *Caspase-8* KD-Induced Enhancement of RA Signaling

Procaspase-8 and FADD inhibit not only activation of RA signaling but also necroptosis mediated by RIPK1, RIPK3 and MLKL. We then investigated whether these necroptosis-inducing genes were involved in the enhancement of RA signaling. Intriguingly, the marked enhancement of RA-induced differentiation was cancelled by down-regulated expression of *Ripk3* using sh*Ripk3* in Casp8 KD ES cells (Figures 2C,D). The enhancement of RA-specific genes expressions and RARE-dependent transcription of luciferase were also cancelled by down-regulating expression of *Ripk3* in *Casp8* KD P19 cells (Figures 2E-G). In addition, *Ripk1* was also shown to be involved in the evident enhancement of RA signaling in *Casp8* KD ES cells (Figure 2 - figure supplement 5A,B). By contrast, MLKL was not involved in the enhancement of RA signaling in *Casp8* KD ES cells (Figure 2 - figure supplement 5C,D). We then asked whether the enhancement of RA signaling was mediated by the kinase activities of RIPK1 and RIPK3, which are required for the induction of necroptosis. In sh*Ripk3*-expressing *Casp8* KD P19 cells, *Casp8* KD-induced enhancement of RA signaling was restored by exogenous expression of shRNA-resistant cDNAs encoding Wt *Ripk3* and a kinase-negative mutant, K51A *Ripk3* (Zhang et al., 2009) (Figures 2E,F). Treatment of *Casp8* KD P19 cells with a kinase inhibitor of RIPK1, Nec-1 (Degterev et al., 2005), did not inhibit the *Casp8* KD-induced enhancement of RA signaling (Figure 2 - figure supplement 5E). These results indicate that both RIPK1 and RIPK3, but not their kinase activities, play an essential role in the enhancement of RA signaling induced by KD of *Casp8* expression. Collectively, the *Casp8* KD-induced enhancement of RA signaling was shown to be dependent on RIPK1 and RIPK3 but independent of their necroptosis-inducing activities.

**Figure 2 - figure supplement 5.**
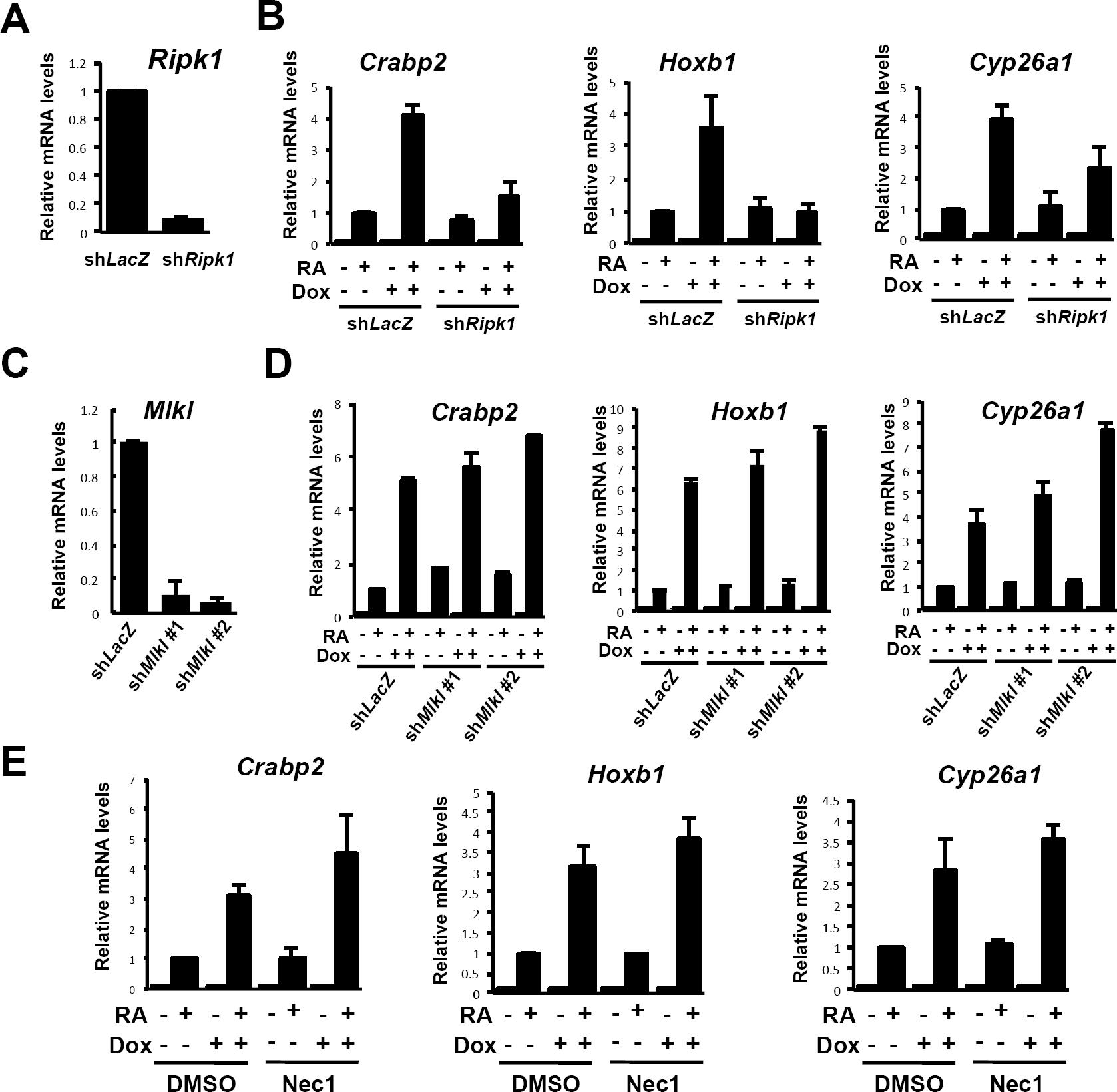
RIPK1 but not MLKL is involved in the activation of RA signaling in the absence of Casp8. (A) Expression levels of Ripk1 were analyzed by qRT-PCR using P19 cells expressing sh*LacZ* or sh*Ripkl.* Representative data are shown as means ± SEM (n=3).
(B) Tet-On sh*Casp8* P19 cells expressing sh*LacZ* or sh*Ripkl* were cultured for 4 days with or without 1 μg/ml Dox and then treated with or without 1 μM RA for 24 h in the presence or absence of Dox. Subsequently, qRT-PCR analysis of RA-induced genes, *Crabp2, Hoxbl* and *Cyp26al,* was performed. Representative data are shown as means ± SEM (n=3).
(C) Expression levels of *Mlkl* were analyzed by qRT-PCR using P19 cells expressing sh*LacZ* or sh*Mlkl.* Two shRNAs targeting different nucleotide sequences in *Mlkl* (sh*Mlkl #1* and sh*Mlkl #2*) were used. Representative data are shown as means ± SEM (n=3).
(D) Tet-On shCasp8 P19 cells expressing s sh*LacZ,* sh*Mlkl #1* or sh*Mlkl #2* were cultured for 4 days with or without 1 μg/ml Dox and then treated with or without 1 μM RA for 24 h in the presence or absence of Dox.. Subsequently, qRT-PCR analysis of RA-induced genes, Crabp2, Hoxbl and Cyp26al, was carried out. Representative data are shown as means ± SEM (n=3).
(E) qRT-PCR analysis of RA-induced genes, *Crabp2, Hoxb1,* and *Cyp26a1,* was performed using Dox (1 μg/ml)-trcatcd or -untreated Tet-On sh*Casp8* P19 cells cultured with or without 1 μM RA for 24 h in the presence of DMSO or 30 μM Nec-1. Representative data are shown as means ± SEM (n=3).

### Knockdown of *Caspase-8* Expression Sensitizes Cells in EBs to RA-Induced Necroptosis

Two-day RA treatment of EBs derived from *Casp8* KD ES cells were found to be smaller than EBs from Tet-On *shCasp8* ES cells treated with Dox but not with RA or those treated with RA but not with Dox (Figures 3A,B). We then analyzed whether cell death was induced in the EBs by LDH release assay, indicating that cell death was clearly and specifically induced in the RA-treated *Casp8* KD EBs (Figure 3C). Since caspase-8 was reported to suppress necroptosis mediated by RIPK1, RIPK3 and MLKL (Hitomi et al., 2008; He et al., 2009, Zhang et al., 2009; Wang et al., 2014), we analyzed the effect of necrostatin-1 (Nec-1), a kinase inhibitor for RIPK, on the RA-induced cell death in *Casp8* KD EBs. The cell death was inhibited by treatment with Nec-1 (Figures 3A-C), which was shown to be unable to inhibit the marked enhancement of RA signaling in *Casp8* KD ES cells (Figure 2 - figure supplement 5E). These results indicated that the mode of death was necroptosis. RA-induced necroptosis in *Casp8* KD EBs was inhibited by exogenous expression of Wt Casp8 and CS Casp8 but not by DE Casp8 (Figure 3 - figure supplement 1), indicating that protease activity of procaspase-8 regulates RA-induced necroptosis as well as RA signaling. Moreover, KD of *Ripk3* expression, which could inhibit both necroptosis and the marked enhancement of RA signaling, strongly inhibited RA-induced necroptosis in *Casp8* KD ES cells (Figure 3C). To summarize thus far, induced KD of *Casp8* expression in EBs derived form TetR-sh*Casp8* ES cells markedly and simultaneously sensitized them to RA-induced cell differentiation and necroptosis.

**Figure 3.**
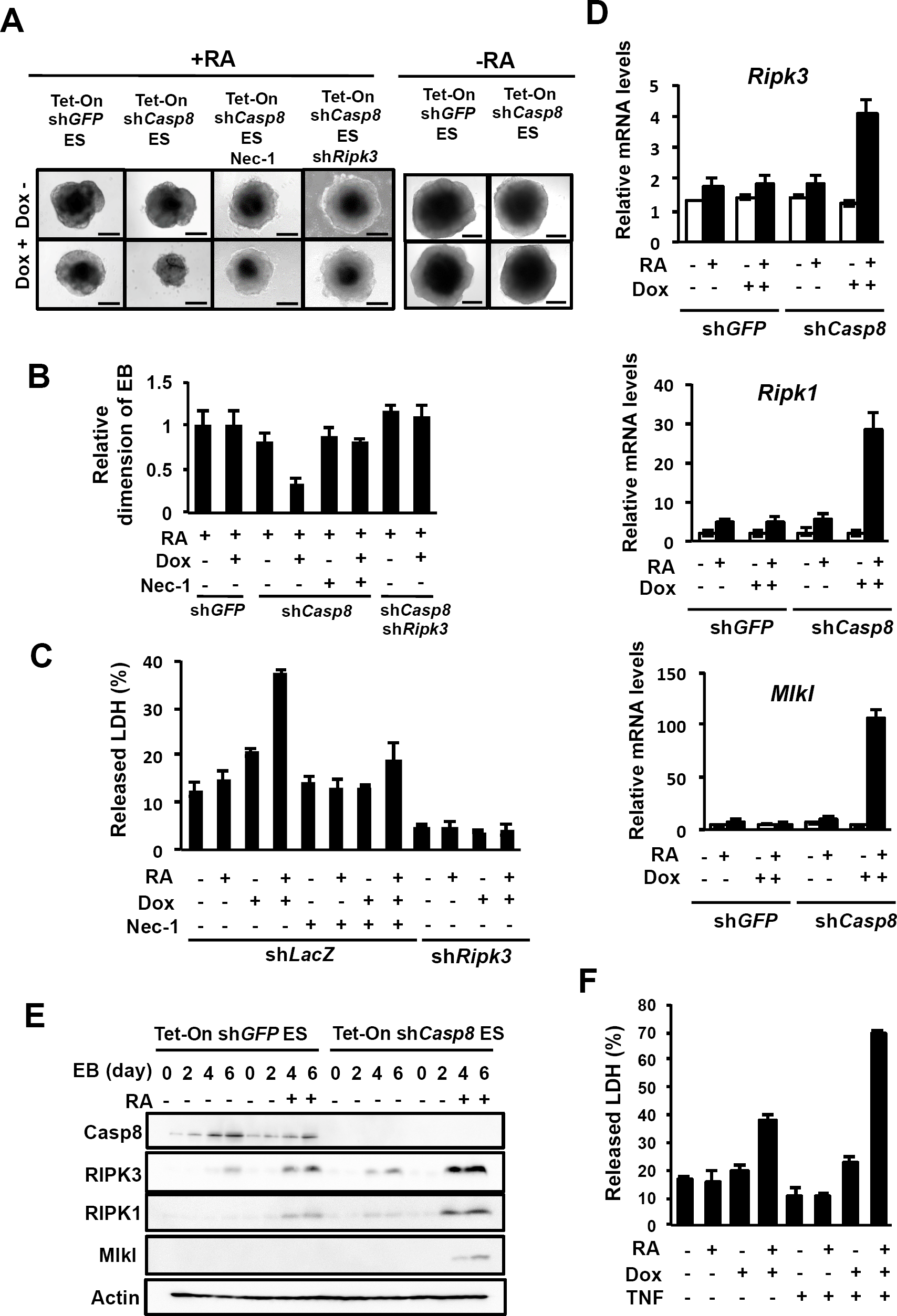
Knockdown of *Casp8* in ES cells enhances RA-induced necroptosis. (A) Dox (1 μg/ml)-treated or -untreated Tet-On sh*Casp8* or Tet-On sh*GFP* ES cells with or without expression of sh*Ripk3* were analyzed by phase-contrast microscopy after 4 days formation of EBs. EBs were treated with or without 1 μM RA and 30 μM Nec-1 for last 2 days. Scale bar, 200 μm.
(B) Relative diameters of EBs generated as described in (A) were quantified. Representative data are shown as means ± SEM (n=10).
(C) Cell death was quantified by a lactate dehydrogenase (LDH) release assay after 4 days of formation of EBs derived from Tet-On sh*GFP* ES cells or Tet-On sh*Casp8* ES cells with or without expression of sh*Ripk3*. EBs were treated with or without 1 μM RA and 30 μM Nec-1 for last 2 days. Representative data are shown as means ± SEM (n=3).
(D) qRT-PCR analysis of *Ripk1, Ripk3* and *Mlkl* were performed using Tet-On sh*GFP* and Tet-On sh*Casp8* ES cells after 6 days formation of EBs in the presence or absence of 1 μg/ml Dox. EBs were cultured with or without 1 μM RA for last 4 days. Representative data are shown as means ± SEM (n=3).
(E) Western blot analysis of *Casp8, Ripk1, Ripk3* and *Mlkl* were performed using Tet-On sh*GFP* and Tet-On sh*Casp8* ES cells at the indicated times after formation of EBs in the presence or absence of 1 μg/ml Dox. EBs were cultured with or without 1 μM RA after 2 days formation of EBs. Actin was detected as a control.

To investigate the mechanism of RA-induced necroptosis in *Casp8* KD EBs, expression levels of *Ripk1, Ripk3* and *Mlkl* were quantified in EBs derived from *Casp8* KD ES cells in the presence or absence of RA. During RA-induced differentiation, expression levels of *Ripk1, Ripk3* and *Mlkl* in control ES cells were slightly increased, but their RA-induced expression levels were remarkably up-regulated by KD of *Casp8* expression in both mRNA and protein levels (Figures 3D,E). In addition, tumor necrosis factor (TNF), which induces necroptosis in various types of cells, enhanced necroptosis in RA-treated *Casp8* KD EBs, but not in RA-untreated *Casp8* KD EBs (Figure 3F). Thus, the enhancement of RA signaling by *Casp8* KD should sensitize *Casp8* KD cells in EBs to necroptosis induced by TNF through the up-regulation of *Ripkl, Ripk3* and *Mlkl* expression.

**Figure 3- figure supplement 1.**
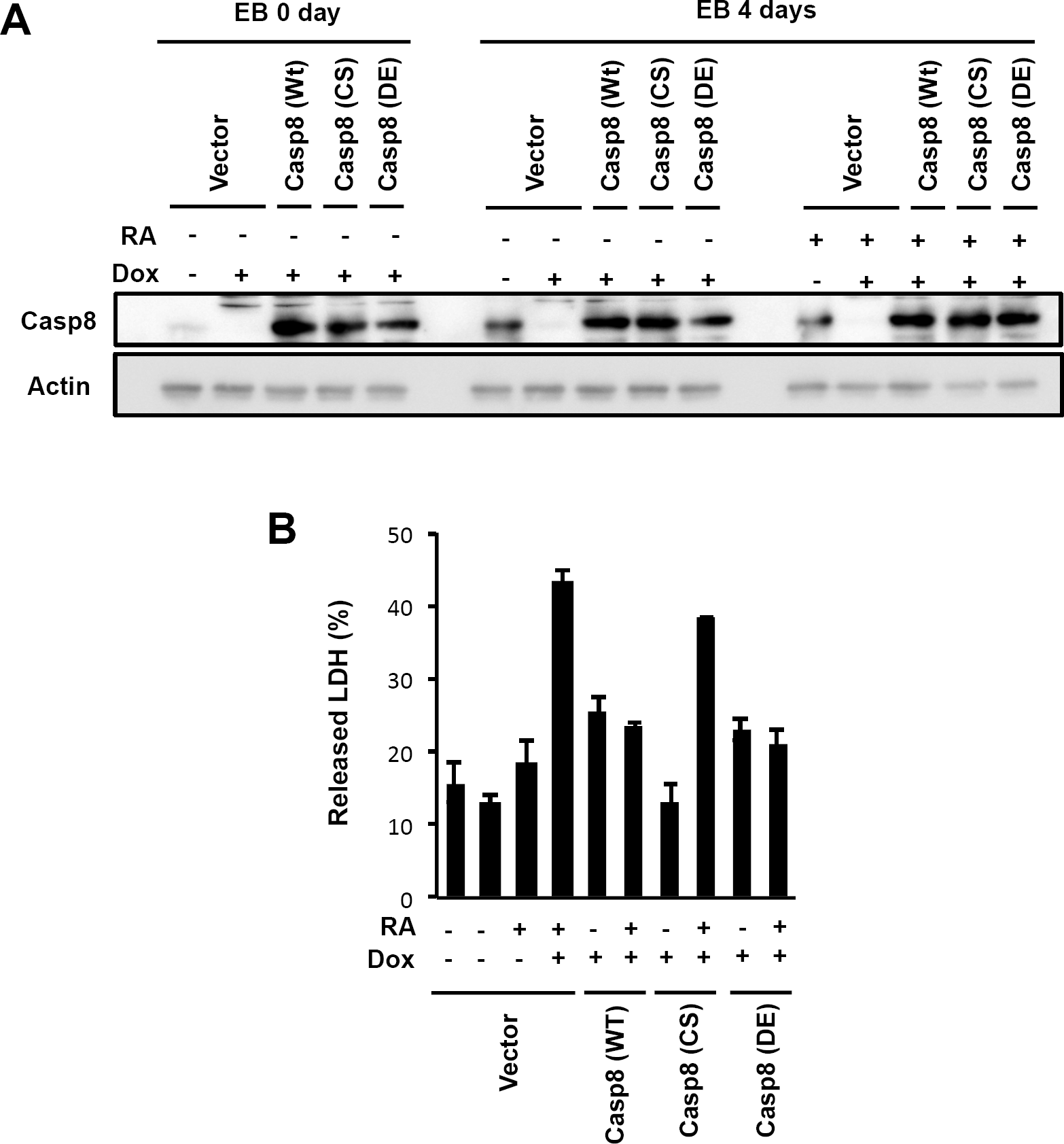
Protease activity of procaspase-8 regulates the RA-induced necroptosis, related to Figure 3. (A) Western blot analysis with an anti-caspase-8 antibody was performed using Tet-On sh*Casp8* ES cells expressing Wt, CS and DE *Casp8* cDNA constructs at the indicated times after formation of EBs in the presence or absence of 1 mg/ml Dox. EBs were cultured with or without 1 μM RA after 2 days formation of EBs. Actin was detected as a control.
(B) Cell death was quantified by an LDH release assay after 4 days of formation of EBs derived from Tet-On sh*Casp8* ES cells expressing WT, CS and DE *Casp8* cDNA constructs. EBs were treated with or without 1 μM RA for last 2 days. Representative data are shown as means ± SEM (n=3).

### KD of *Caspase-8* Expression Induces Nuclear Translocation of RIPK1 and RIPK3 to Form a Complex with RARs

To explore the molecular mechanism underlying the *casoase-8* KD-induced enhancement of RA signaling, we examined the relationship between RIPKs and RA signaling. Since RIPK1 and RIPK3 are localized in the cytoplasm under normal conditions and RA binds to RARs in the nucleus, we at first analyzed whether RIPK1 and RIPK3 can translocate into the nucleus. Treatment of Casp8-expressing P19 cells with Leptomycin B (LMB), an inhibitor of nuclear export of proteins (Yang et al., 2004), converted subcellular localization of RIPK3 from cytoplasm to both cytoplasm and nucleus (Figure 4A). LMB treatment induced not only nuclear localization of RIPK3 but also promotion of RA signaling in a RIPK3-dependent manner (Figure 4B), suggesting that RIPK3 in the nucleus enhances RA signaling. In addition, KD of *Casp8* expression as well as treatment with LMB converted subcellular localization of RIPK3 from cytoplasm to both cytoplasm and nucleus (Figures 4C,D). Taken together, caspase-8 suppresses nuclear localization of RIPK3, and intranuclear RIPK3 may play an important role in the enhanced activation of RA signaling in the absence of caspase-8.

**Figure 4.**
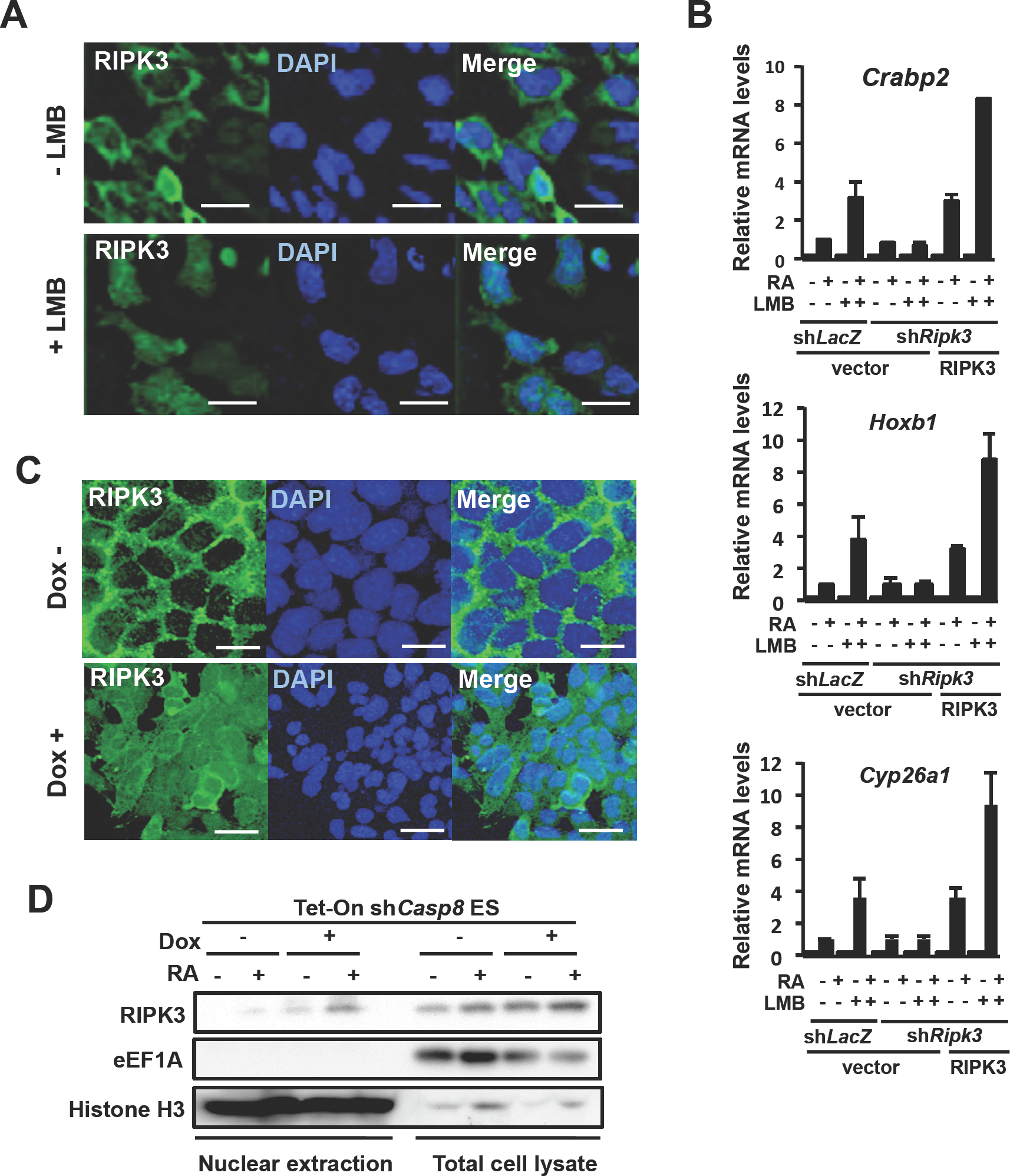
KD of *Casp8* expression induces nuclear translocation of RIPK1 and RIPK3 to enhance RA signaling. (A) P19 cells expressing sh*Ripk3* and sh*Ripk3*-resistant 3xFlag-Ripk3 were treated with or without 1 μM RA and 2 ng/ml LMB for 24 h, and then subcellular localization of 3xFlag-RIPK3 was analyzed by fluorescence microscopy after staining with DAPI. Scale bar, 20 μm.
(B) P19 cells expressing sh*Ripk3* and sh*Ripk3*-resistant 3xFlag-*Ripk3* were subjected to qRT-PCR analysis of RA-induced genes, *Crabp2, Hoxb1* and *Cyp26a1,* after treatment with or without 1 μM RA and 2 ng/ml LMB for 24 h. vector, an empty vector. Representative data are shown as means ± SEM (n=3).
(C) Tet-On sh*Casp8* P19 cells expressing sh*Ripk3* and sh*Ripk3*-resistant 3xFlag-Wt *Ripk3* were cultured with or without1 μg/ml Dox for 4 days in the presence or absence of Dox. Subsequently, subcellular localization of 3xFlag-RIPK3 was analyzed by fluorescence microscopy using anti-Flag antibody after staining with DAPI. Scale bar, 20 μm.
(D) Western blot analysis of endogenous RIPK3 from nuclear fractions and total cell lysates was carried out using Tet-On sh*Casp8* ES cells cultured with or without 1 μg /ml Dox for 5 days. Cells were treated with or without 1 μM RA for last 24 h. Cytoplasmic eEF1A1 and nuclear Histone H3 were simultaneously analyzed.

Subcellular localization of exogenously expressed mCherry-RIPK3 in P19 cells was also converted from cytoplasm to both cytoplasm and nucleus by overexpression of RARα, which is a nuclear protein (Figure 5A). Overexpressed RARα seemed to retain RIPK3 in nucleus. We then analyzed the interaction of RARα with RIPK1 and RIPK3. In co-immunoprecipitation experiments, exogenously expressed RIPK3 interacted with exogenously expressed RARα in HEK293T cell extracts (Figure 5B), and immunoprecipitation of exogenous Flag-tagged RIPK3 with anti-Flag antibody co-precipitated endogenous RARα in *Casp8* KD ES cells (Figure 5C). Importantly, endogenous RIPK1 and RIPK3 were co-immunoprecipitated with endogenous RARα with an anti-RAR antibody from lysates of *Casp8* KD ES cells (Figure 5D). Moreover, the interactions of RARα to both RIPK1 and RIPK3 were shown to be enhanced by treatment with RA. Thus, KD of *Casp8* expression induced nuclear localization of RIPK3, and RIPK1 and RIPK3 interacted with RARα in the nucleus might activate RA signaling.

**Figure 5.**
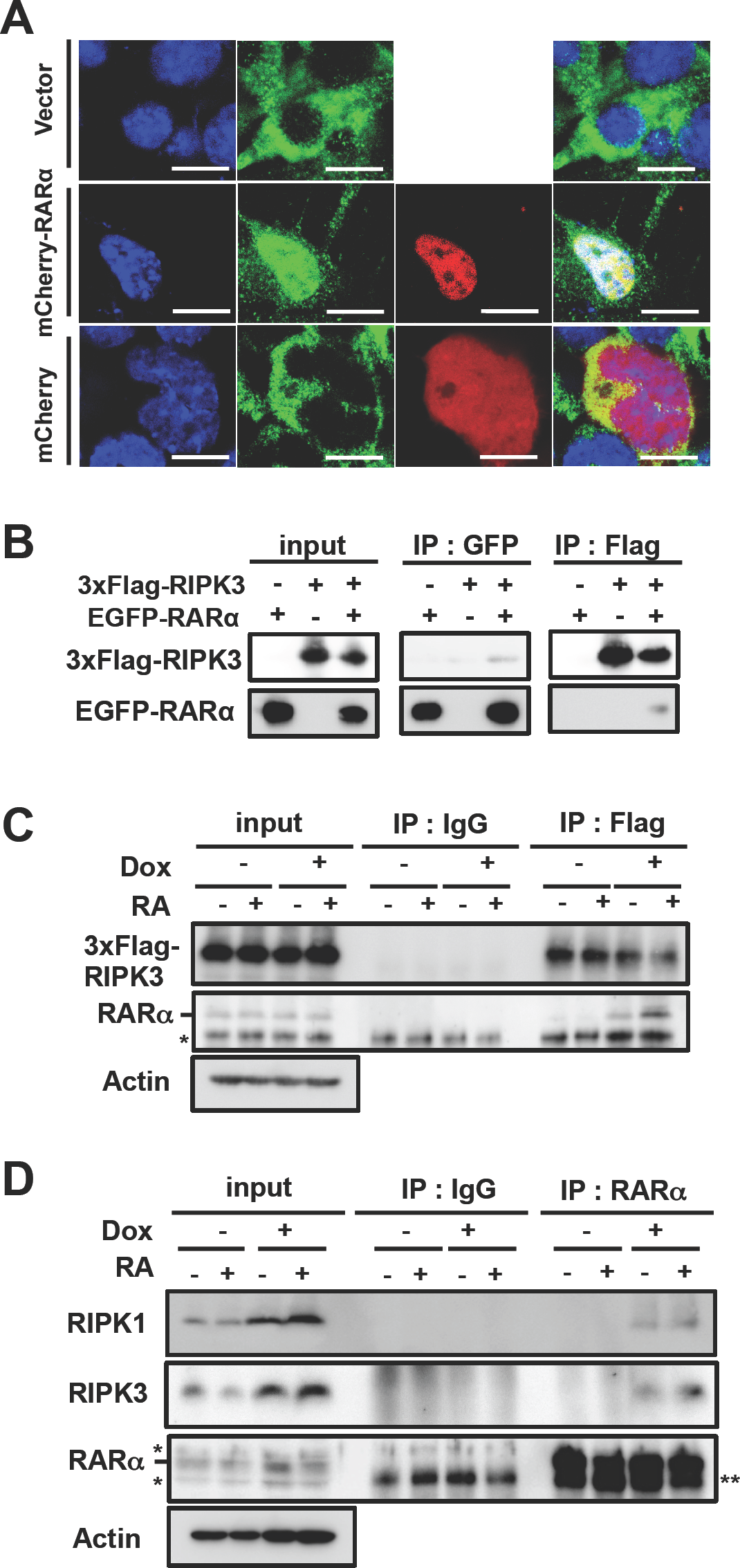
RIPK1 and RIPK3 interact with RARα. (A) P19 cells expressing sh*Ripk3* and sh*Ripk3*-resistant 3xFlag-Ripk3 were transfected with an expression vector encoding mCherry-RARα or mCherry, and cultured for 48h. Then, subcellular localization of mCherry-RARα or mCherry was analyzed by fluorescence microscopy after staining with DAPI. Scale bar, 20 μm.
(B) Lysates of HEC293T cells transiently expressing 3xFlag-tagged RIPK3 and/or EGFP-RARα were subjected to immunoprecipitation with anti-Flag antibody or anti-GFP antibody, and analyzed by western blotting with anti-Flag antibody or anti-GFP antibody. Total cell lysates (input) were also analyzed.
(C and D) Western blot analysis was carried out for immunoprecipitates (IP) with control IgG and ant-Flag Ab from Tet-On sh*Casp8* ES cells expressing 3xFlag-Wt *Ripk3* with control IgG or anti-RARα from Tet-On sh*Casp8* ES cells (C), or IP with control IgG or anti-RARα from Tet-On sh*Casp8* ES cells (D).). Total cell lysates (Input) were also analyzed. *: nonspecific bands, **: IgG-derived bands.

In co-immunoprecipitation experiments, exogenously expressed RIPK3 interacted with exogenously expressed RARα in HEK293T cell extracts (Figure 5B), and co-immunoprecipitation analyses using various deletion mutants of RARα and RIPK3 indicated that RIPK3 and RARα interacted through the ligand-binding domain (LBD) of RARα and the protein kinase domain (PKD) of RIPK3 (Figure 5 - figure supplement 1). Since RIPK3 was reported to directly bind to RIPK1 through its RHIM domain, RIPK3 would be able to function as an adaptor protein through binding to RIPK1 and RARα *via* its RHIM and kinase domains, respectively.

**Figure 5 - figure supplement 1.**
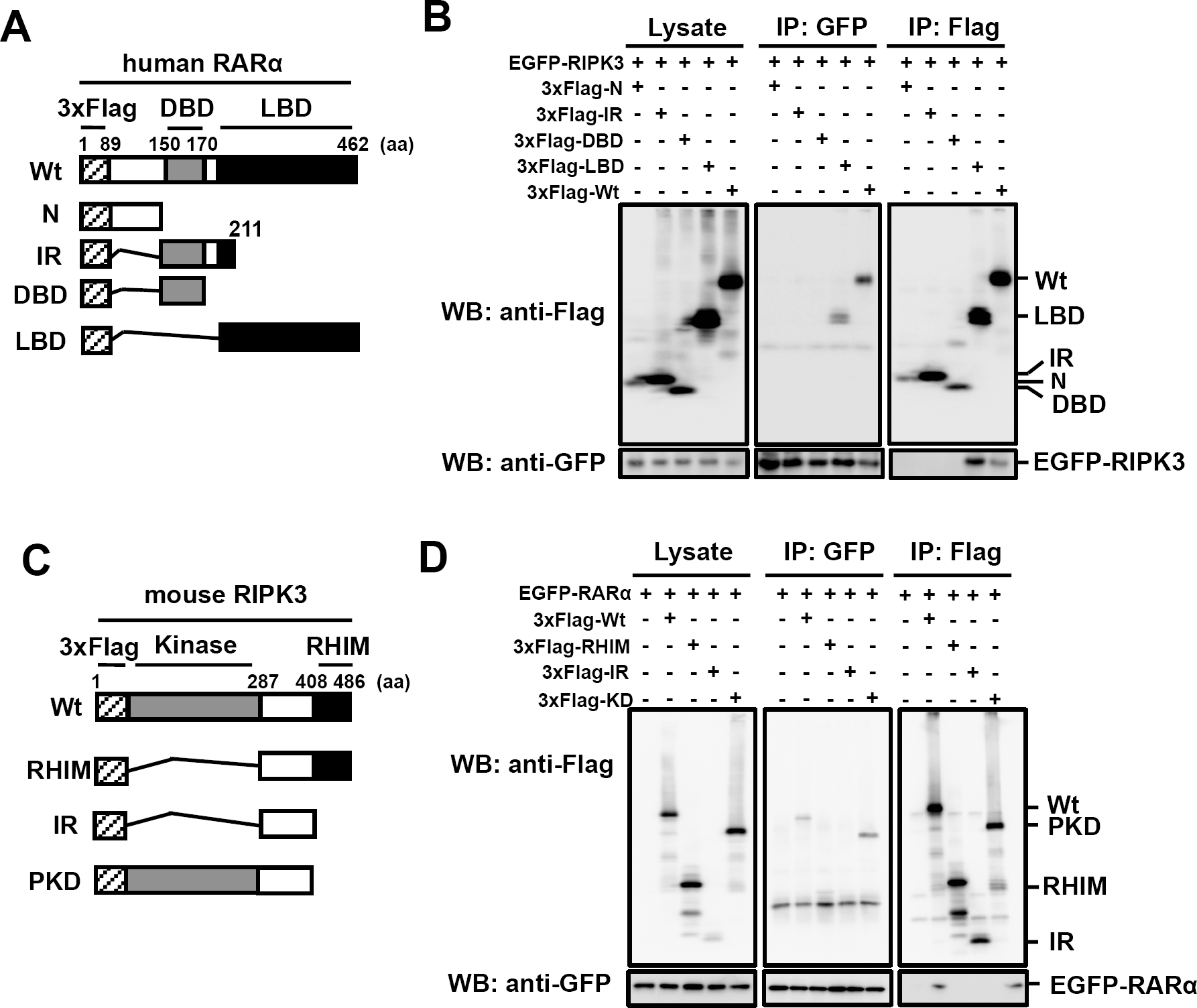
The ligand-binding domain (LBD) of RARα interacts with the protein kinase domain (PKD) of RIPK3, related to Figure 5. (A) Graphical overview of 3xFlag-tagged full-length human RARα (Wt) and its various deletion mutants. The numbers of amino acid residues in RARα are indicated. N, N-terminal domain of RARα; IR, intermediate region of RARα; DBD, DNA-binding domain of RARα; and LBD, ligand binding domain of RARα.
(B) Lysates of HEC293T cells transiently expressing EGFP-RIPK3 and various deletion mutants of 3xFlag-RARα were subjected to immunoprecipitation with anti-Flag or anti-GFP antibodies, and IP were analyzed by western blotting with anti-Flag or anti-GFP antibodies. Total cell lysates (Lysate) were also analyzed.
(C) A graphical overview of 3xFlag-tagged full-length mouse RIPK3 (Wt) and its various deletion mutants. Relevant numbers of amino acid residues in RIPK3 are indicated. RHIM, RIPK homotypic interaction motif (RHIM) domain; IR, intermediate region; and PKD, protein kinase domain.
(D) HEC293T cells transiently expressing EGFP-RARα or various deletion mutants of 3xFlag-RIPK3 were subjected to immunoprecipitation with anti-Flag or anti-GFP antibodies, and analyzed by western blotting with anti-Flag or anti-GFP antibodies. Total cell lysates (Lysate) were also analyzed.

### The Complex of RIPK1, RIPK3 and RARs Works in The Coactivator Complex to Enhance RA-Dependent Transcription

To clarify whether the complex of RIPK1, RIPK3 and RARs functions in RA-dependent transcription, chromatin immunoprecipitation (ChIP) analysis was carried out in *Casp8* KD ES cells. RA treatment was indicated to notably enhance binding of endogenous RIPK1 to *RARE* of an RA-inducible gene, *Rarb,* specifically in the absence of *Casp8* expression (Figure 6A). RA treatment also clearly enhanced binding of exogenously expressed Wt and a kinase-negative mutant, K51A RIPK3 to *RARE* of an RA-inducible gene, *Cyp26a1,* in the absence of endogenous *Ripk3* and *Casp8* expressions (Figure 6B). Furthermore, KD of Casp8 significantly enhanced RA-induced biding of RARs to *RARE* dependently on *Ripk3* expression but independently on the kinase activity of RIPK3 (Figure 6C). Taken together, RARs, bound to RIPK1 and RIPK3 in the nucleus of *Casp8* KD cells, showed much stronger binding activity to RARE in the presence of RA than RARs without RIPK1 and RIPK3.

*Caspase-8* KD-induced enhancement of RA signaling was shown to be dependent upon RXRα, TDG, p300 and CBP (Figure 6 - figure supplement 1), all of which were reported to form a transcriptional coactivator complex with RAR on *RAREs* and to enhance RA signaling (Um et al., 1998; Lee et al., 2009; Cortellino et al., 2011). Collectively, all the results suggest that RIPK1 and RIPK3 form a coactivator complex with RAR/RXR, TDG, p300 and CBP in the nucleus of *Casp8* KD cells, RA treatment induces tight binding of the RIPK1/RIPK3-containing coactivator complex onto *RAREs,* and RA signaling is subsequently enhanced.

**Figure 6.**
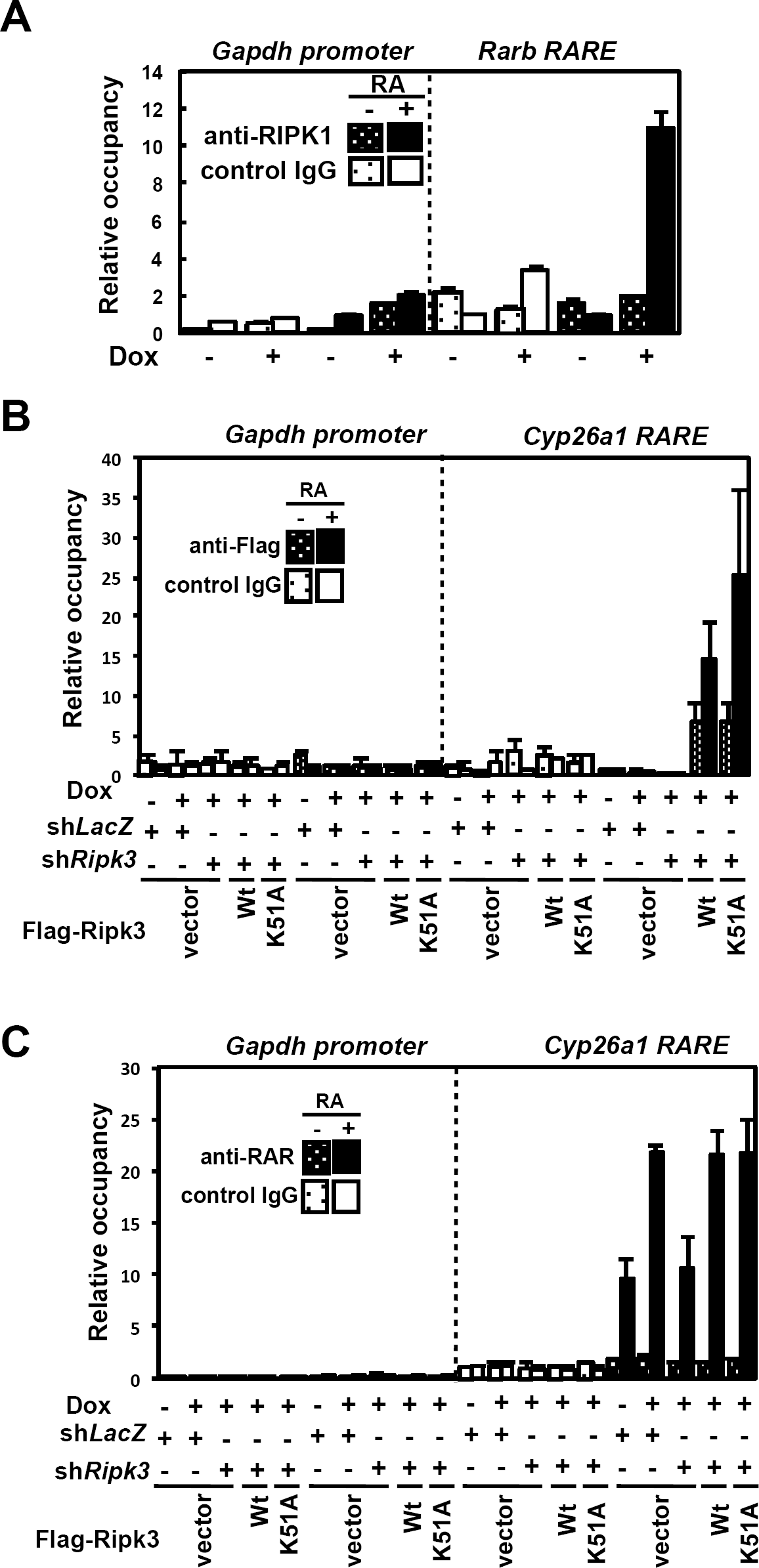
RA treatment notably enhances binding of not only RARs but also RIPK1 and RIPK3 to *RAREs* of RA-inducible genes in the absence of *Casp8* expression. (A) Tet-On sh*Casp8* and sh*GFP* ES cells were cultured with or without 1 μg/ml Dox for 4 days and then treated with or without 1 μM RA for 24 h in the presence or absence of Dox. Subsequently, ChIP analysis for the *Rarb*-specific *RARE* using an anti-RIPK1 antibody was carried out. Representative data are shown as means ± SEM (n=3).
(B and C) Tet-On sh*Casp8* P19 cells expressing sh*LacZ* or sh*Ripk3* together with or without sh*Ripk3*-resistant 3xFlag-Wt or K51A Ripk3 (kinase-negative form of RIPK3) were treated with or without 1 μM RA for 24 h in the presence or absence of 1 μg/ml Dox, and subjected to ChIP analysis using anti-Flag antibody or control IgG (B), and anti-RAR antibody or control IgG (C). vector, an empty vector. Representative data are shown as means ± SEM (n=3).

**Figure 6 - figure supplement 1.**
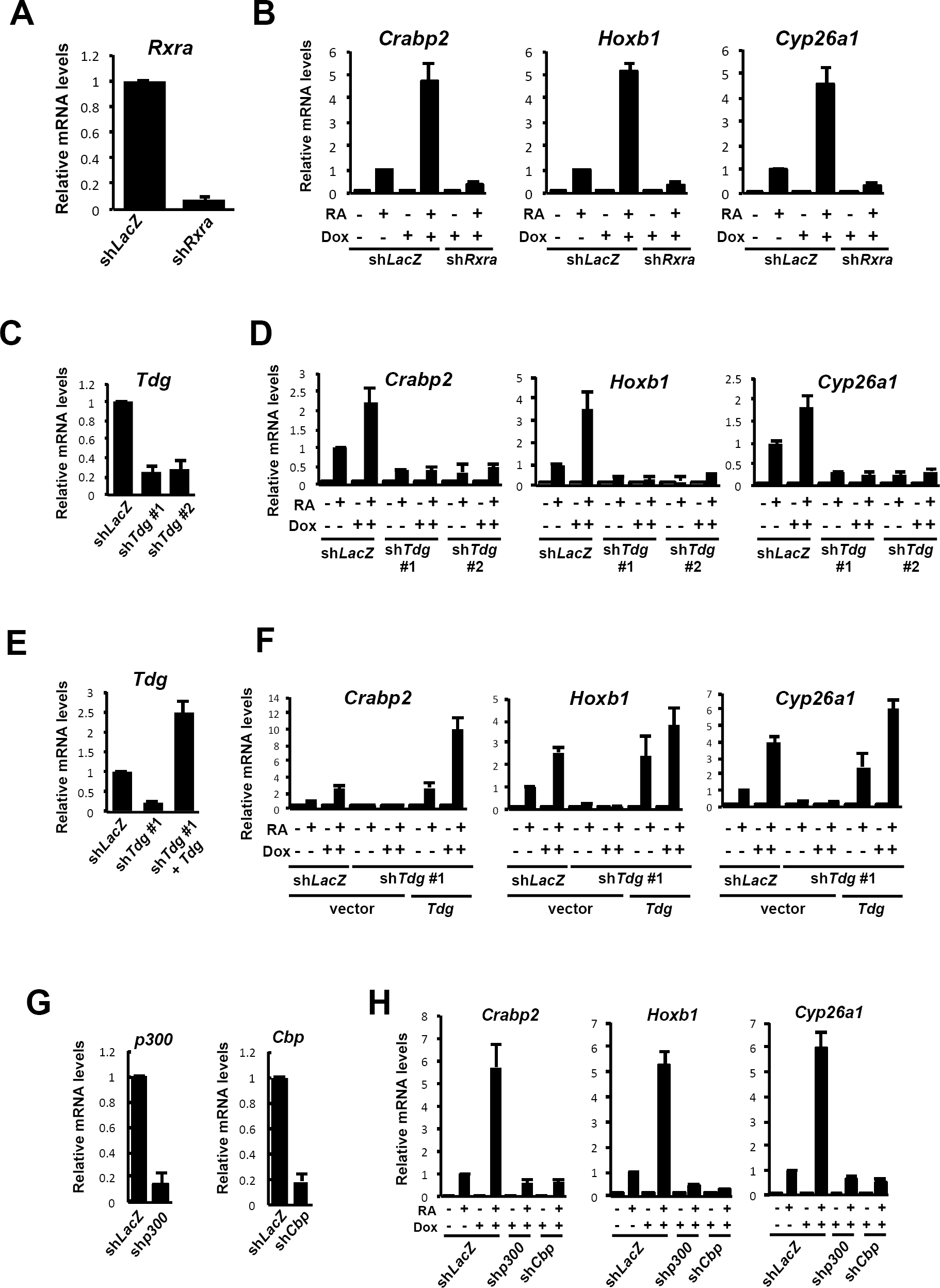
RXRα, TDG, p300 and CBP are required for the marked enhancement of RA signalling in the absence of *Casp8* expression, related to Figure 6. (A) P19 cells expressing sh*LacZ* or sh*Rxra* were subjected to qRT-PCR analysis for *Rxra.* Representative data are shown as means ± SEM (n=3).
(B) P19 cells expressing sh*LacZ* or sh*Rxra* were cultured for 4 days with or without 1 μg/ml Dox and then treated with or without 1 μM RA for 24 h. Subsequently, qRT-PCR analysis of RA-induced genes, *Crabp2, Hoxb1* and *Cyp26a1,* was carried out. Representative data are shown as means ± SEM (n=3).
(C) Expression levels of *Tdg* were analyzed by qRT-PCR using P19 cells expressing sh*LacZ* or sh*Tgd.* Two shRNAs targeting different nucleotide sequences in *Tdg* (sh*Tdg #1* and sh*Tdg #2*) were used. Representative data are shown as means ± SEM (n=3).
(D) P19 cells expressing sh*LacZ*, sh*Tdg #1* or sh*Tdg #2* were cultured for 4 days with or without 1 μg/ml Dox and then treated with or without 1 μM RA for 24 h. Subsequently, qRT-PCR analysis of RA-induced genes, *Crabp2, Hoxb1* and *Cyp26a1,* was carried out. Representative data are shown as means ± SEM (n=3).
(E) P19 cells expressing sh*LacZ* or sh*Tdg #1* together with or without Wt *Tdg* were subjected to qRT-PCR analysis for *Tdg.* Representative data are shown as means ± SEM (n=3).
(F) P19 cells expressing sh*LacZ* or sh*Tdg #1* together with or without Wt *Tdg* were cultured with or without 1 μM RA for 24 h in the presence or absence of 1 μg/ml Dox, and then subjected to qRT-PCR analysis for RA-inducible genes. vector, an empty vector. Representative data are shown as means ± SEM (n=3).
(G) P19 cells expressing sh*LacZ* or *shp300,* and sh*LacZ* or sh*Cbp* were subjected to qRT-PCR analysis for *p300* and *Cbp,* respectively. Representative data are shown as means ± SEM (n=3).
(H) P19 cells expressing sh*LacZ*, sh*p300* or sh*Cbp* were cultured for 4 days with or without 1 μg/ml Dox and then treated with or without 1 μM RA for 24 h. Subsequently, qRT-PCR analysis of RA-induced genes, *Crabp2, Hoxb1* and *Cyp26a1,* was carried out. Representative data are shown as means ± SEM (n=3).

### RA Signaling Is Enhanced in *Caspase-8^-/-^* Mouse Embryos

*Caspase-8*-deficient (*Casp8^-/-^*) mouse embryos die at E11.5 with associated abnormal yolk sac vascularization, heart development and neural tube formation (Sakamaki et al., 2002). In mouse embryonic fibroblasts (MEFs) from *Casp8^-/-^* mice (Figure 7A), RA-induced gene expression as well as RARE-dependent transcription of luciferase was enhanced, compared to *Casp8^+/+^* MEFs (Figures 7B,C). We also detected up-regulated transcription of RA-induced genes in *Casp8^-/-^* whole embryos at E10.5 by qRT-PCR (Figure 7D) and *in situ* hybridization analyses (Figure 7E). The up-regulation was more prominent in the embryo at E11.5. *In situ* hybridization analyses revealed that expression patterns of RA-inducible genes in *Casp8^-/-^* embryos were essentially the same as those of Wt embryos. Taken together, RA signaling was supposed to be enhanced in *Casp8^-/-^* embryos as well as *Casp8* KD cells.

**Figure 7.**
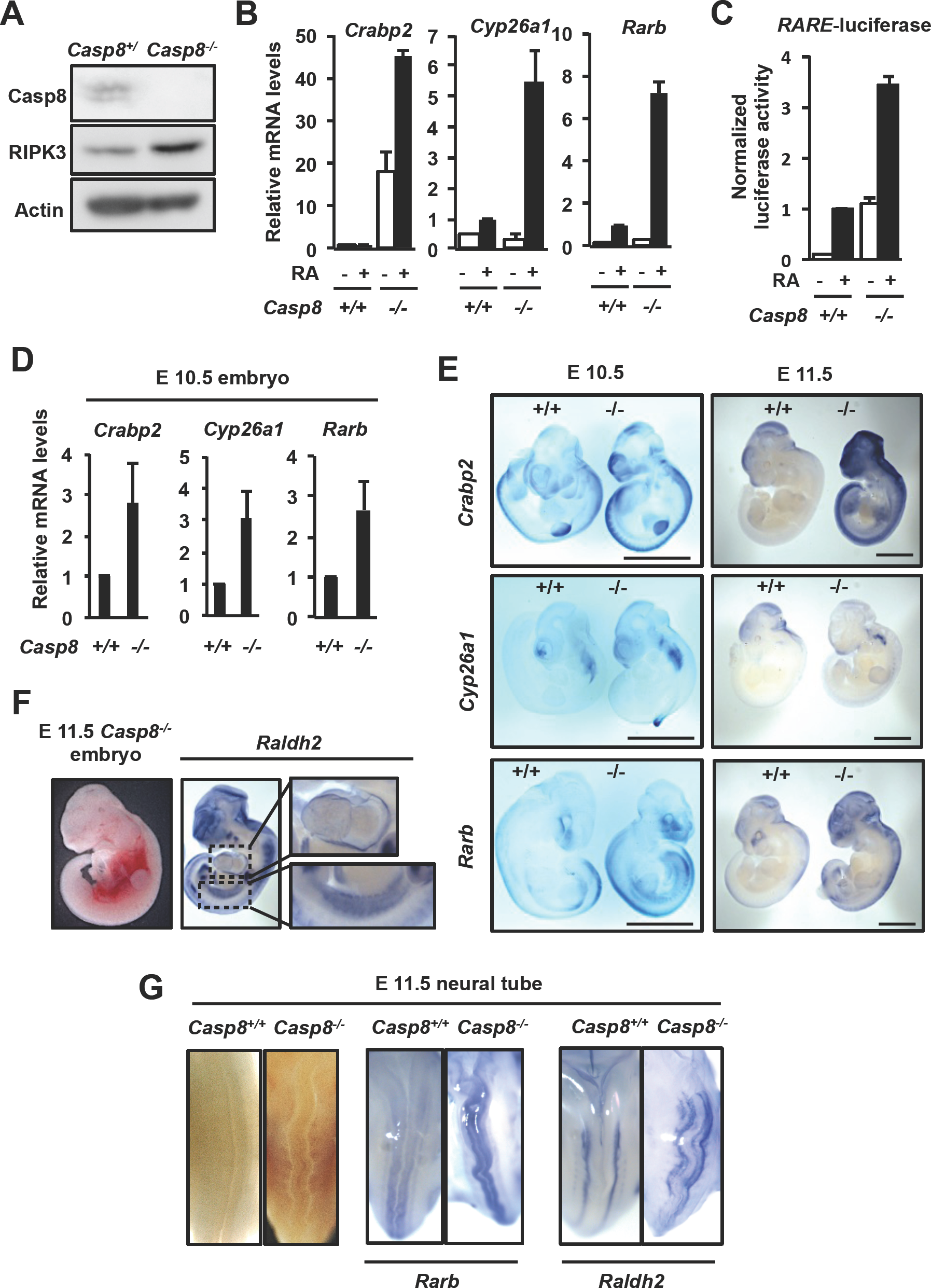
RA signaling is enhanced in *Casp8^-/-^* mouse embryos. (A) Immortalized MEFs derived from Wt *(Casp8^+/+^)* and *Casp8^-/-^* embryos were subjected to western blot analysis for caspase-8 and RIPK3.
(B) qRT-PCR analysis of RA-induced genes, *Crabp2, Cyp26a1* and *Rarb,* was carried out in *Casp8^+/+^* and *Casp8^-/-^* MEFs after treatment with or without 1 μM RA for 24 h. Representative data are shown as means ± SEM (n=3).
(C) Dual-luciferase reporter analysis of *RARE* was carried out using RA-treated *Casp8^+/+^* and *Casp8^+/-^* MEFs. Representative data are shown as means ± SEM (n=3).
(D) qRT-PCR analysis of RA-inducible genes, *Crabp2, Cyp26a1* and *Rarb,* was performed in E10.5 Wt *(Casp8^+/+^)* and *Casp8^-/-^* littermates. RNA was extracted from whole embryos.
(E) Expressions of RA-inducible genes, *Crabp2, Cyp26a1* and *Rarb,* in E10.5 or E11.5 Wt *(Casp8^+/+^)* and *Casp8^-/-^* littermates were detected by whole-mount in situ hybridization analysis. Scale bar, 2 mm.
(F and G) Whole-mount *in situ* hybridization analysis of *Rarb* and *Raldh2* was carried out using E11.5 *Casp8^-/-^* embryos. Views of heart and AGM (F), and neural tube (G) were shown.

In addition, a prominent RA response was observed in tissues of *Casp8^-/-^* embryos, heart, aorta-gonad-mesonephros (AGM) and neural tube, all of which were reported to be abnormal in *Casp8^-/-^* embryos (Figures 7F,G). These results seem to show the possibility that enhancement of RA signaling is at least in part involved in the embryonic letality of *Casp8^-/-^* mice, although the embryonic letality of *Casp8^-/-^* mice has been shown to be due to excess necroptosis (Dillon et al., 2016). A chemical inhibitor of RA signalling, BMS493 (le Maire et al., 2010), was intraperitoneally injected into the pregnant *Casp8* heterozygous (*Casp8^+/-^*) female mice intercrossed with *Casp8^+/-^* male mice (Figure 8A). The up-regulated expression of RA-specific genes in *Casp8^-/-^* embryos was partly but significantly suppressed by treatmetent with BMS493 (Figure 8B). BMS493-treated Casp8-deficient embryos were observed to be viable even at E11.5, and the characteristic abnormal phenotypes of yolk sac, neural tube and heart in *Casp8*-deficient embryos were also rescued by injection of BMS493 (Figures 8C-E). It should be noted, however, that BMS493-treated E12.5 Casp8-deficient embryos were not viable. Thus, chemical inhibition of RA signalling, which is enhanced in *Casp8^-/-^* embryos, delays, but does not completely inhibit, embryonic lethality of *Casp8*-deficient embryos.

**Figure 8.**
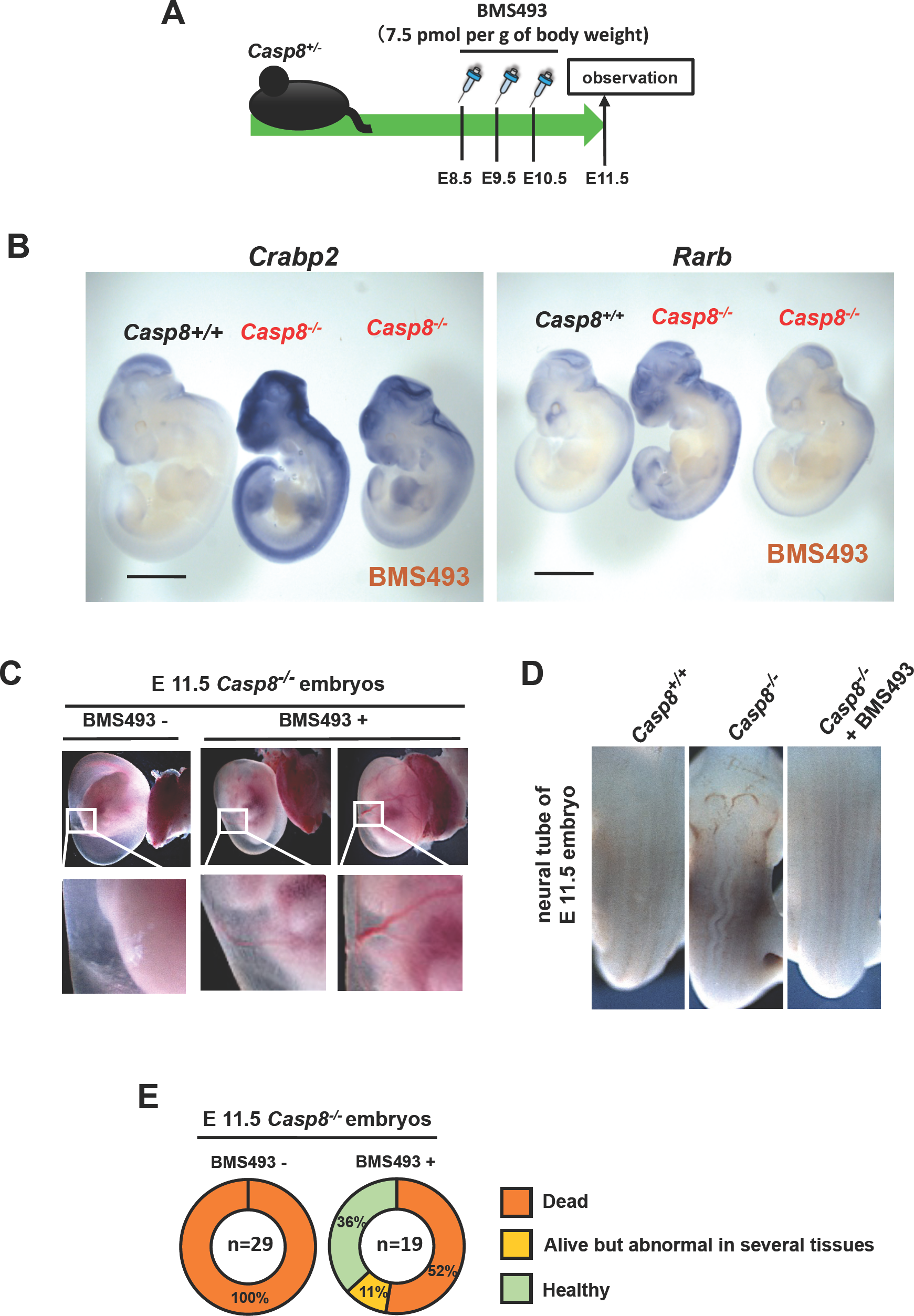
Marked enhancement of RA signaling in the absence of Casp8 is correlated with abnormality of *Casp8^-/-^* embryos. (A) Experimental design of intra-peritoneal injection of BMS493 into pregnant mice.
(B) Whole-mount in situ hybridization analysis of RA-inducible genes, *Crabp2* and *Rarb,* was carried out for E11.5 Wt *(Casp8^+/+^)* and *Casp8^-/-^* littermates, and E11.5 *Casp8^-/-^* embryos from pregnant *Casp8^+/-^* mice treated with BMS493 (scale bar, 2 mm).
(C) Views of yolk sac of E11.5 *Casp8^-/-^* embryos treated with or without BMS493.
(D) Views of neural tube of E11.5 *Casp8^+/+^* and *Casp8^-/-^* embryos, and *Casp8^-/-^* embryo treated with BMS493.
(E) Quantification of viability and abnormalities of E11.5 *Casp8^-/-^* embryos treated with or without BMS493.

## Discussion

The enhancement of RA-specific genes expression in *Casp8* KD ES cells was significantly induced by not only 1 μM RA (Figure 2A), but also 100 and 10 nM RA (Figure2 - figure supplement 1A). Since the concentration of RA in mouse embryo tissues was reported to be approximately 25 nM on average (Horton,. 1995), caspase-8 was supposed to suppress evident activation of RA signaling in a physiological condition. Indeed, up-regulated transcription of RA-induced genes was detected in *Casp8^-/-^* whole embryos by qRT-PCR (Figure 7D) and *in situ* hybridization analyses (Figure 7E). Furthermore, expressions of sh*Casp8*-resistant Wt *Casp8* significantly inhibited the evident activation of RA signaling by *Casp8* KD ES cells (Figure2 - figure supplement 4)). Based on all the *in vitro* and *in vivo* findings in this manuscript, suppression of caspase-8 or FADD expression, or caspase-8 activity generally enhances RA signaling/RARE-dependent transcription through nuclear translocation of RIPK1 and RIPK3 followed by their association with RARs in a coactivator complex (Figure 9).

**Figure 9.**
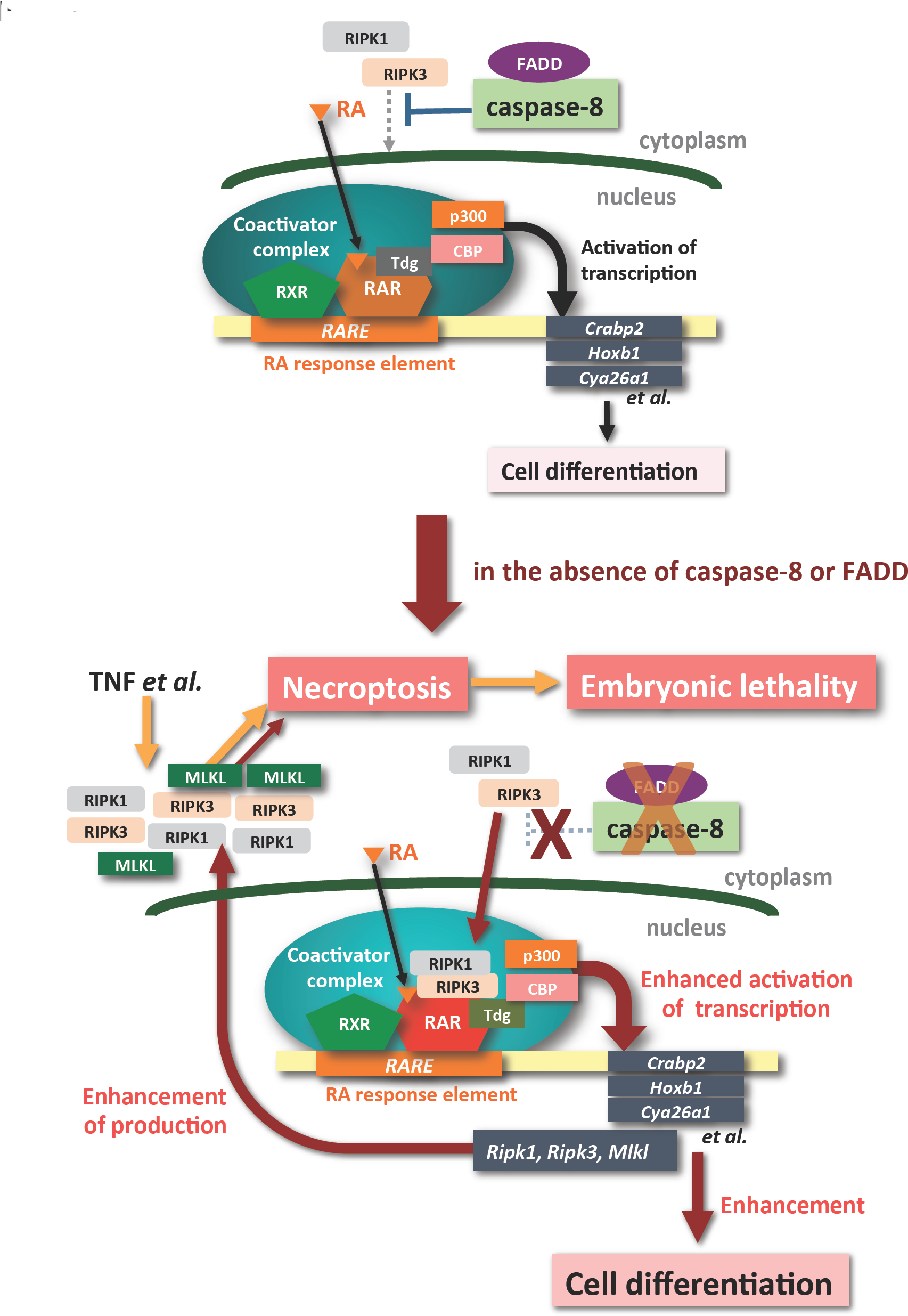
A model of the RA signalling in the presence and absence of caspase-8. Brown directional lines indicate signaling pathways induced and/or enhanced by the absence of caspase-8 activity/expression or FADD expression.

Induced KD of *Casp8* expression in EBs derived form TetR-sh*Casp8* ES cells was shown to sensitize the cells to RA-induced necroptosis (Figure 3C). RA-induced necroptosis was observed in EBs derived from *Casp8* KD ES and P19 cells, but not in either *Casp8* KD ES and P19 cells before EB formation, SK-N-SH cells or HL60 cells. RA-induced necroptosis seems to be induced in only EB or its related early embryo. In fact, the cell lines, which were insensitive to RA-induced necroptosis, were sensitized to apoptosis by strong stimulation with RA (data not shown). Yet-to-be-defined mechanism(s) in EBs might contribute the conversion of response to RA, from apoptosis to necroptosis. Necroptosis was induced in RA-treated *Casp8* KD EBs through enhanced expression of *Ripk1, Ripk3* and *Mlkl* (Figure 3D). RARs were reported to be able to directly bind to a RARE-like sequence in the promoter region of *Ripk1* in RA-treated ES cells (Moutier et al., 2012). The RA-induced up-regulation of *Ripk1* expression in *Casp8* KD EBs might be directly mediated by RARs during RA-induced differentiation of cells in EBs. However, it has not been determined yet whether the RA-induced up-regulation of *Ripk3* and *Mlkl* are directly mediated RARs or not.

*Casp8^-/-^* embryos die at E10.5-E11.5 with associated abnormal yolk sac vascularization, heart development and neural tube formation ( Varfolomeev et al., 1998; Sakamaki et al., 2002). The embryonic lethality of *Casp8^-/-^* embryos is rescued by not only depletion of *Ripk3* (Kaiser et al., 2011) but also knock-in of kinase-negative K51A *Ripk3* (Mandal et al., 2014), indicating that kinase activity of RIPK3, which is indispensable for induction of necroptosis, is required for embryonic lethality of *Casp8^-/-^* embryos. In contrast, RIPK3 is, but kinase activity of RIPK3 is not, required for the enhancement of RA signaling in *Casp8* KD ES cells (Figures 2G). In addtion, *Casp8^-/-^ Mlkl^-/-^* mice were recently reported to be viable and to mature into fertile adults (Alvarez-Diaz et al., 2016). Thus, MLKL and kinase activity of RIPK3, both of which are required for the induction of necroptosis but not for the enhancemnt of RA signaling in *Casp8* KD cells, are indispensable for embryonic letality of *Casp8^-/-^*embryos. Taken togetger, the embryonic letality of *Casp8^-/-^* mice is due to excess necroptosis (Dillon et al., 2016), but not due to the enhancement of RA-induced differentiation. Interestingly, *Casp8^-/-^ Mlkl^-/-^* mice succumbed significantly earlier than *Casp8^-/-^ Ripk3^-/-^* mice due to a more rapid onset of severe lymphadenopathy and autoimmune pathology (Alvarez-Diaz et al., 2016), suggesting that biological activity other than necroptosis-inducing activity of RIPK3 is involved in the rapid onset of severe autoimmunity in *Casp8^-/-^ Mlkl^-/-^* mice. It would be interesting to analyze whether the RIPK3-dependent enhancement of RA signaling is implicated in the rapid onset of severe autoimmunity in *Casp8^-/-^ Mlkl^-/-^* mice.

Elimination of TNFR1 from *Casp8^-/-^* embryos was shown to delay embryonic lethality from E10.5 until E16.5 (Dillon et al., 2014), indicating that TNFR1-mediated necroptosis plays an essential role in the embryonic letality of *Casp8^-/-^* mice around E10.5. Hence RA-induced necroptosis should not be required for the embryonic letality of *Casp8^-/-^* mice around E10.5. However, TNF was shown to induce necroptosis in RA-treated but not RA-untreated *Casp8* KD EBs (Figure 3F), suggesting that enhancement of RA signaling contributes sensitization of cells in *Casp8^-/-^* embryo to TNF-induced necroptosis. In addition, chemical inhibition of RA signaling delayed embryonic letality of *Casp8^-/-^* mice until E12.5 (Figures 8E). Taken together, we suppose that *ripk1, ripk3* and *mlkl* expressions are enhanced by RA in *Casp8^-/-^* embryos, and their increased expressions might be partly involved in the embryonic letality of *Casp8^-/-^* mice around E10.5 through enhancing the sensitivity to TNF-induced necroptosis (Figure 9).

Caspase-8 and RIPK were shown to regulate sensitivity to RA signalling in not only mouse ES cells but also human cancer cell lines (Figure2 - figure supplement 3). Intriguingly, acute promyelocytic leukemia (APL) was reported to be dramatically improved by co-treatment with RA and a caspase inhibitor when compared with treatment with only RA (Ablain et al., 2014). Taken together, our study contributes to a deeper understanding of the nexus of caspase-8 and RA signaling in cell death, cell differentiation, development, stem cell fate, ontogenesis, and the fate of cancer cells.

## Materials and methods

### Mice

C57BL/6 mice were purchased from CLEA Japan. *Casp8^-/-^* mice were generated as described previously (Sakamaki et al., 2002). C57BL/6 *Casp8^-/-^* mice were bred and maintained in specific pathogen-free conditions. All experiments in this study were performed according to the guidelines for animal treatment at the institute of Laboratory Animals (Kyoto University).

### Cell culture

TT2 mouse ES cells were maintained on mitomycin C-treated MEFs in Dulbecco’s modified Eagle’s medium (DMEM, Gibco) supplemented with 1% fetal bovine serum (JRH Bioscience), 10% knockout serum replacement (KSR, Gibco), 2 mM L-glutamine, 0.1 mM β-mercaptoethanol, 0.1 mM non-essential amino acids (Gibco), 1 mM sodium pyruvate (Gibco) and 1000 U/ml LIF (CHEMICOM). MEFs, P19 cells, SK-N-SH cells and HEC293T cells were cultured in Dulbecco’s modified Eagle’s medium (DMEM, Nacalai Tesque Inc.) supplemented with 10% fetal bovine serum (Sigma), 100 U/ml penicillin and 100 μg/ml streptomycin (Nacalai Tesque Inc.). HL60 cells, a kind gift from K. Inaba (Kyoto University), were cultured in suspension culture in RPMI 1640 (Nacalai Tesque Inc.). All cells were cultured at 37°C in 5% CO_2_.

### Plasmids, lentiviral expression vectors, and shRNA expression system

p*RARE3*-Luciferase and an expression vector carrying human RARα cDNA were kind gifts from A. Kakizuka (Kyoto University). Lentiviral vectors, originally provided by H. Miyoshi (RIKEN), were prepared as described previously^31^. For expression of mouse *Casp8* with sh*Casp8*-resistant silent mutations, the corresponding cDNA fragment after site-directed mutagenesis was subcloned into pCSII-PGK-MCS-IRES-Hyg. For expression of mouse *Tdg* and mouse *Ripk3* with sh*Ripk3*-resistant silent mutations, their corresponding cDNA fragments were subcloned into pCSII-PGK-3xFlag-MCS-IRES-Hyg. To generate stably shRNA-expressing cells, we utilized lentivirus vectors, pCSII-U6-MCS and pCSII-U6-MCS-puro (kind gifts from Y. Satoh and M. Matsuoka, Kyoto University). shRNA-encoding DNA oligonucleotides were inserted into these vectors. To achieve the specific knockdown of mouse *Casp8* or *Fadd,* the tetracycline inducible shRNA expression system (Tet-On shRNA system) with lentivirus based vectors (pCSII-EF-TetR-IRES-puro and pCSII-U6tet-sh*Casp8* or sh*Fadd*-PGK-neo) was utilized in TT2 mouse ES and P19 cells as previously described (Kobayashi and Yonehara, 2009; Kikuchi et al., 2012).

**Table 1.**
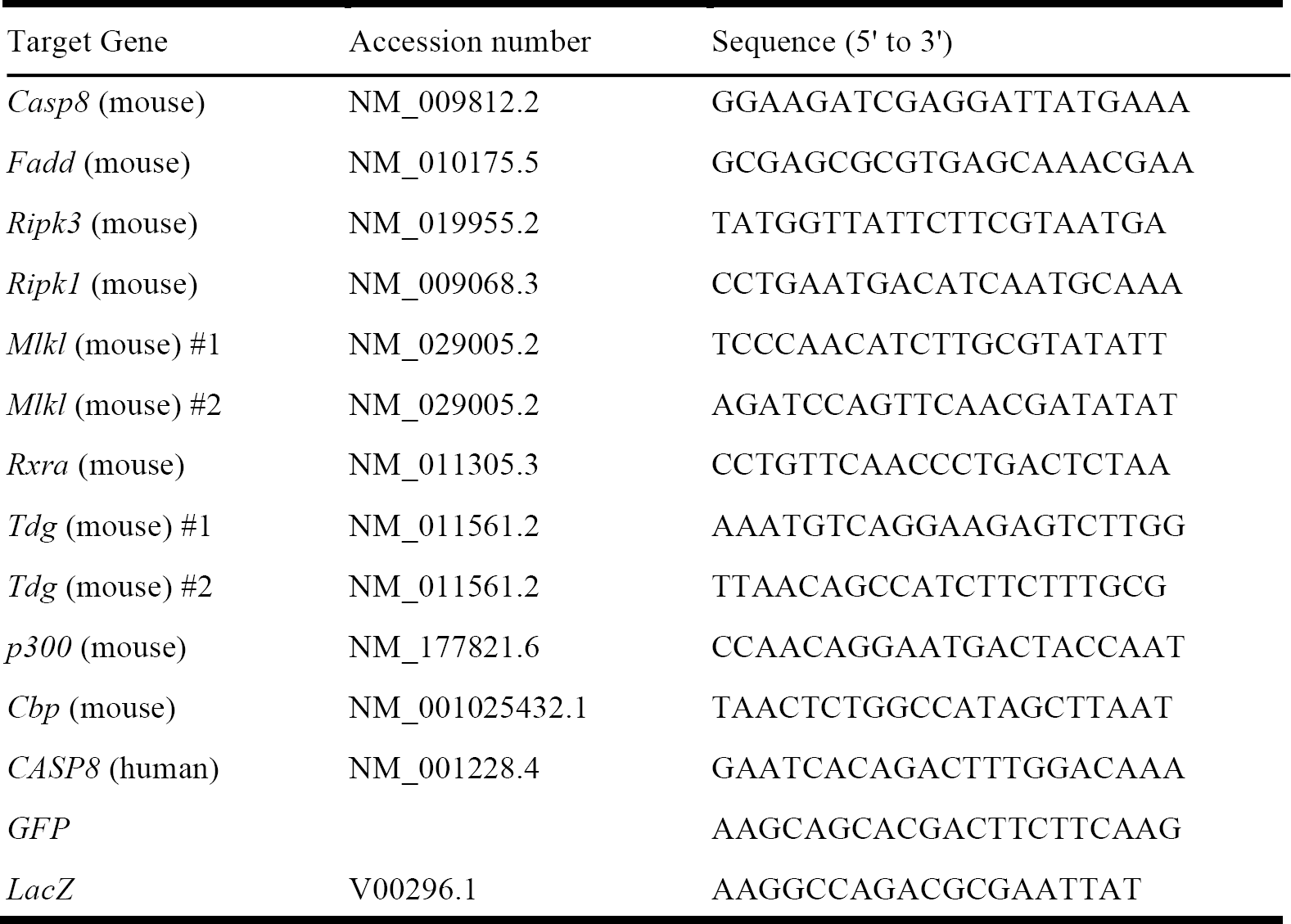
The following primer sequences of shRNA were used:

### RA treatment for quantification of RA-induced genes

Cells were treated with 1 μM, 100 nM, 10nM or 1nM RA (Sigma) for 24 h, and then expression levels of RA-induced genes were quantified.. In the case of cells with a Tet-On shRNA system, cells were cultured with or without 1 (μg/ml Dox for 4 days, and then treated with or without 1 RA for 24 h in the presence of Dox. To analyze expression of genes directly induced by RA, cells were treated with RA for only 24 h.

### Differentiation of ES cells and P19 cells through EB formation

After Tet-On sh*Casp8* or sh*Fadd* ES cells were cultured with or without 1 μg/ml Dox for 2 days, single-cell suspensions were prepared by treatment with trypsin-EDTA (Nacalai Tesque Inc.). To form EBs, 3 × 10^6^ cells were seeded per well in low-cell-adhesion 96-well plates (Thermo SCIENTIFIC) in Glasgow’s Minimum Essential Medium (GMEM, Gibco) supplemented with 10% knockout serum replacement (KSR, Gibco), 2mM L-glutamine, 0.1 mM μ-mercaptoethanol, 0.1 mM non-essential amino acids (Gibco) and 1 mM sodium pyruvate (Gibco) (ES differentiation medium) in the presence of Dox. Two days after seeding, medium was changed to ES differentiation medium supplemented with or without 1 μM RA. After 2 days cultivation, formed EBs were transferred to collagen type I-coated chamber slides (Becton Dickinson), cultured for 4 days in ES differentiation medium supplemented with or without 1 μM RA (RA treatment was for 6 days in all.), and subjected to immunohistochemical analysis. To induce significant differentiation of cells through EB formation, 6 days treatment with RA was necessary.

For RA-induced neural differentiation of Tet-On sh*Casp8* P19 cells, cells were treated with or without 1 μg/ml Dox for 4 days, and single-cell suspensions were prepared by treatment with trypsin-EDTA (Nacalai Tesque Inc.). To form EBs, 1 × 10^6^ cells were seeded per 10 cm non-treated dish (IWAKI) in DMEM (Nacalai Tesque Inc.) supplemented with 10% fetal bovine serum (Sigma), 100 U/ml penicillin, and 100 μg/ml streptomycin (Nacalai Tesque Inc.), and cultured for 2-6 days with or without 1 μM RA.

## LDH release assay

After Tet-On sh*Casp8* or *shCasp8/Ripk3* ES cells were cultured with or without 1 μg/ml Dox for 2 days, single-cell suspensions were prepared by treatment with trypsin-EDTA (Nacalai Tesque Inc.). To form EBs, 1.6 × 10^5^ cells were seeded per well in non-treated 6-well plates (IWAKI) in ES differentiation medium in the presence of Dox. Two days after seeding, the medium was changed to ES differentiation medium supplemented with or without 1 μM RA and 1 μg/ml Dox. To inhibit necroptosis, cells were cultured with 30 μM Nec1 (Enzo Life Science) thereafter. After a further 2-day cultivation with or without RA, Dox and Nec-1, the LDH release assay was performed using a Cytotoxicity Detection Kit^PLUS^ (Roche) in accordance with manufacturer’s instructions. At least 3 biological experiments were carried out and data are presented as means ± s.d.

### Western blot analysis and immunoprecipitation

For western blot analysis, cells were lysed in ice-cold lysis buffer (20 mM Tris-HCl, pH7.4, with 10% glycerol, 1% Triton X-100, 0.5% Nonidet P-40, 150 mM NaCl, and 1 mM EDTA) containing a protease inhibitor cocktail (Nacalai Tesque Inc.). Cell lysates were resolved by sodium dodecyl sulfate-polyacrylamide gel electrophoresis (SDS-PAGE) and analyzed by western blot analysis as described previously (Minamida et al., 2014). For immunoprecipitation, cells were lysed in RIPA buffer (50 mM Tris-HCl, pH 7.5, 150 mM NaCl, 1 mM EDTA, 1% NP-40, and 0.5% sodium deoxycholate containing a protease inhibitor cocktail (Nacalai Tesque Inc.), and immunoprecipitation was performed following standard protocols. Immunoprecipitates were resolved by SDS-PAGE and analyzed by western blotting.

The antibodies used for western blot analyses and immunoprecipitation in this study were anti-mouse caspase-8 (ALX-804-447-C100, Enzo Life Science), anti-RIPK1 (610458, BD Transduction Laboratories), anti-mouse RIPK3 (ADI-905-242-100, Enzo Life Science), anti-human caspase-8 (M032-3, MBL), anti-Flag M2 (F3165, Sigma), anti-MLKL (MABC604, MERCK MILLIPORE), anti-RAR (M-454, Santa Cruz), anti-Histone H3 (601901, BioLegend), and anti-actin (MAB1501, MERCK MILLIPORE).

### Immunocytochemistry and whole-mount in situ hybridization

Cells in chamber slides were fixed with 4% paraformaldehyde (Nacalai Tesque Inc.) in PBS for 15 min and permeabilized by 3 successive treatments with 0.3% Triton X-100 (Nacalai Tesque Inc.) in PBS for 2 h. Cells were treated with primary antibodies for 12 h at 4^o^C, washed 3 times with 0.05% Tween-20 in PBS, and then treated with Alexa Fluor® 488-conjugated anti-mouse IgG (Molecular Probes) for 1 h. Fixed cells were washed 3 times with 0.05% Tween-20 in PBS and mounted with VECTASHIELD Mounting Medium with DAPI (Vector Laboratories). Cells were analyzed under a confocal fluorescent microscope (OLYMPUS). The antibodies for immunocytochemistry used in this study were anti-Flag M2 (F3165, Sigma) and anti-Tuj1 (MAB1637, MECK MILLIPORE). Whole-mount in situ hybridization was performed as described previously (Harima et al., 2013).

### Dual-Luciferase assay

Tet-On sh*GFP* or Tet-On sh*Casp8* TT2 mouse ES cells transfected with p*RARE3*-Luciferase and pTK-*Renilla* luciferase were cultured with or without 1 μg/ml Dox for 5 days and then treated with or without 1 μM RA for 24 h. The Dual-Luciferase assay was performed using a dual-luciferase assay kit (Promega) in accordance with the manufacturer’s instructions. At least 3 biological experiments were carried out and data are presented as means ± s.d.

### qRT-PCR

Total RNA was extracted using Sepasol^®^-RNA Super G (Nacalai Tesque Inc.) according to the manufacturer’s instructions. The reverse transcription (RT) reaction was performed using a ReverTra Ace^®^ qRT-PCR Master Mix (TOYOBO) according to the manufacturer’s instructions. RT products were analyzed using a THUNDERBIRD^®^ qPCR Mix (TOYOBO) and the StepOne real-time PCR system (Applied Biosystems) with the primer sets listed bellow according to the manufacturer’s instructions. The expression level of each mRNA was normalized to that of mouse or human *GAPDH.* At least 3 biological experiments were carried out and data are presented as means ± s.d.

The following primer sequences for qRT-PCR were used:

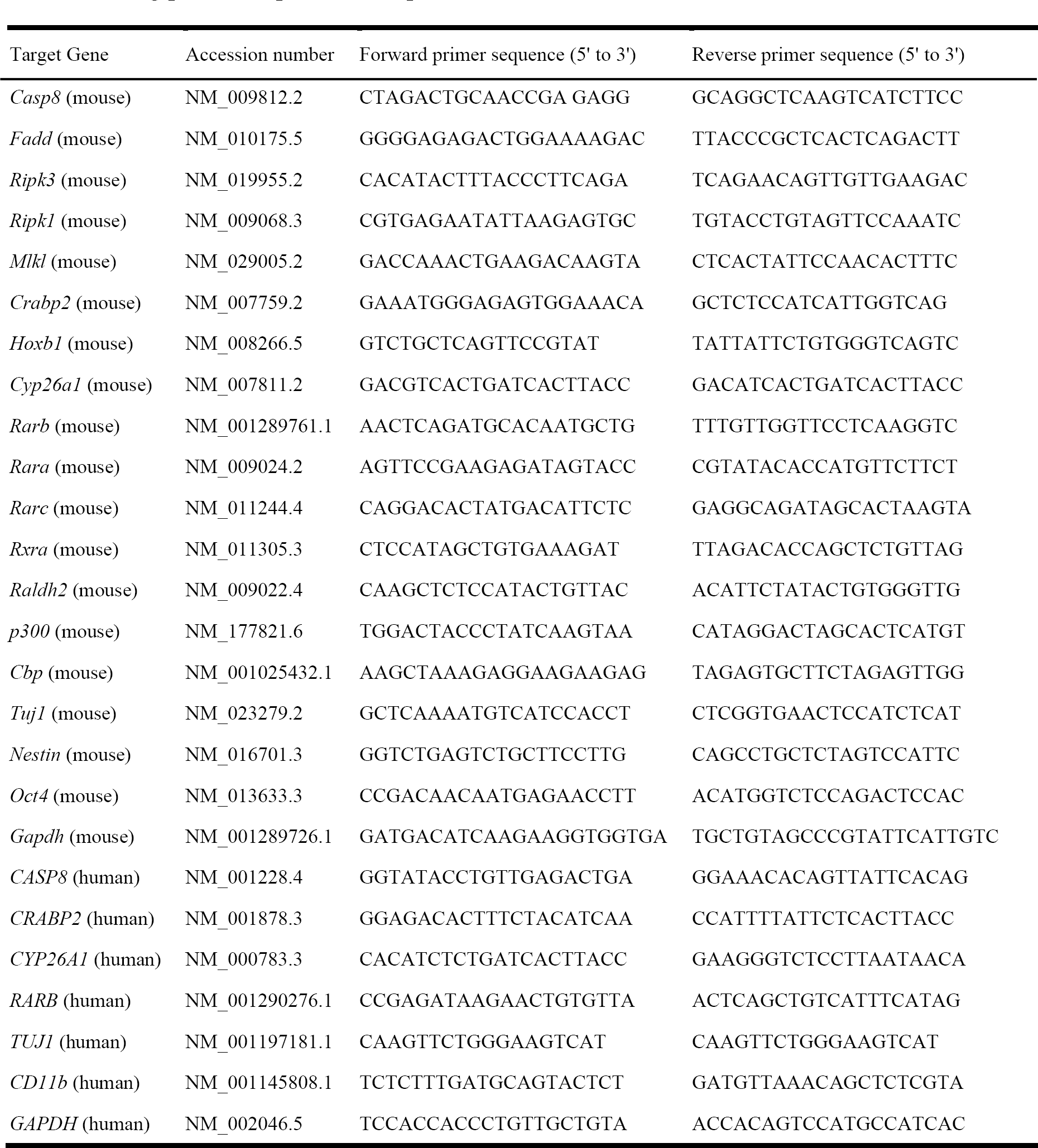

### Flow cytometric analysis

HL60 cells expressing sh*LacZ* or sh*Casp8* were cultured with or without 1 μM RA for 3 days and then stained with FITC-conjugated anti-CD11b antibody (eBioscience) for 30 min. Flow cytometric analysis was performed with a FACS canto2 (BD Biosciences).

### ChIP analysis

ChIP analyses were performed as previously described (Lee et al., 2006). In brief, quantitative PCR was performed using a THUNDERBIRD^®^ qPCR Mix (TOYOBO) and the StepOne real-time PCR system (Applied Biosystems) with the primer listed bellow. The antibodies for ChIP analysis used in this study were anti-Flag M2 (F3165, Sigma), anti-RAR (M-454, Santa Cruz) and anti-RIPK1 (610458, BD Transduction Laboratories). At least 3 biological experiments were carried out and data are presented as means ± s.d.

The following primer sequences for Chip assay were used:

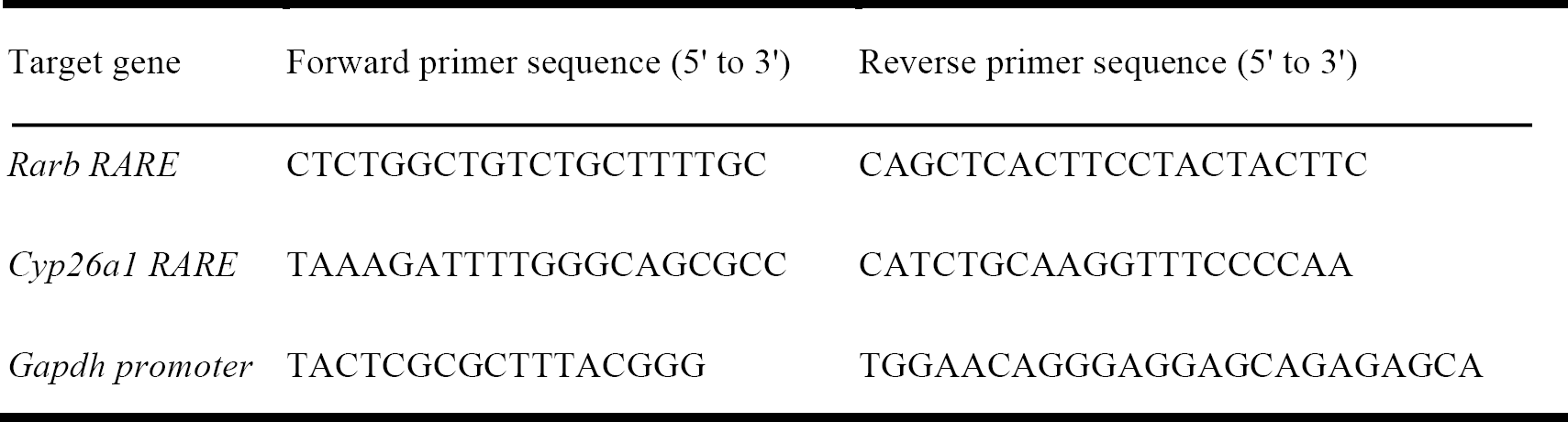

### Rescue experiments of *Casp8^-/-^* embryos using an RA antagonist

BMS493 (Tocris Bioscience) in DMSO (100 mM) was diluted with olive oil to a final concentration of 3μM just before use. BMS493 (2.5 μl/g of body weight; 7.5 pmol/g of body weight) was intraperitoneally injected into pregnant *Casp8^+/-^* female mice intercrossed with *Casp8^+/-^* male mice at E8.5, E9.5 and E10.5 after fertilization, and E11.5 *Casp8^-/-^* embryos were analyzed in comparison with *Casp8^+/+^* littermates.

## Funding

Grant-in-Aid for Scientific Research on Innovative Areas (Homeostatic regulation by various types of cell death) (15H01376) from the Ministry of Education, Culture, Sports, Science and Technology (MEXT) of Japan.

Shin Yonehara

## Acknowledgments

We thank J. A. Hejna for advice and critical reading of the manuscript, and all members of the Yonehara laboratory for helpful discussions. We are grateful to Y. Harima, T. Ohtsuka and R. Kageyama for *in situ* hybridization analyses.

## References

1. Ablain, J., Rice, K., Soilihi, H., de Reynies, A., Minucci, S., and de The, H. (2014). Activation of a promyelocytic leukemia-tumor protein 53 axis underlies acute promyelocytic leukemia cure. Nat. Med. 20, 167–174.

2. Alvarez-Diaz, S., Dillon, C.P., Lalaoui, N., Tanzer, M.C., Rodriguez, D.A., Lin, A., Lebois, M., Hakem, R., Josefsson, E.C., O'Reilly, L.A., et al. (2016). The Pseudokinase MLKL and the Kinase RIPK3 Have Distinct Roles in Autoimmune Disease Caused by Loss of Death-Receptor-Induced Apoptosis. Immunity 45, 513–26.

3. Alnemri, E. (1997). Mammalian cell death proteases: a family of highly conserved aspartate specific cysteine proteases. J. Cell. Biochem. 64, 33–42.

4. Astrom, A., Pettersson, U., Chambon, P., and Voorhees, J.J. (1994). Retinoic acid induction of human cellular retinoic acid-binding protein-II gene transcription is mediated by retinoic acid receptor-retinoid X receptor heterodimers bound to one far upstream retinoic acid-responsive element with 5-base pair spacing. J. Biol. Chem. 269, 22334–22339.

5. Chinnaiyan, A.M., O'Rourke, K., Lane, B.R., and Dixit, V.M. (1997). Interaction of CED-4 with CED-3 and CED-9: a molecular framework for cell death. Science 275, 1122–1126.

6. Cortellino, S., Xu, J., Sannai, M., Moore, R., Caretti, E., Cigliano, A., Le Coz, M., Devarajan, K., Wessels, A., Soprano, D., et al. (2011). Thymine DNA glycosylase is essential for active DNA demethylation by linked deamination-base excision repair. Cell 146, 67–79.

7. Degterev, A., Huang, Z., Boyce, M., Li, Y., Jagtap, P., Mizushima, N., Cuny, G.D., Mitchison, T.J., Moskowitz, M.A., and Yuan, J. (2005). Chemical inhibitor of nonapoptotic cell death with therapeutic potential for ischemic brain injury. Nat. Chem. Biol. 1, 112–119.

8. Dillon, C.P., Weinlich, R., Rodriguez, D.A., Cripps, J.G., Quarato, G., Gurung, P., Verbist, K.C., Brewer, T.L., Llambi, F., et al. (2014) RIPK1 blocks early postnatal lethality mediated by caspase-8 and RIPK3. Cell 157, 1189–1202.

9. Dillon, C.P., Tummers, B., Baran, K., and Green, D.R. (2016). Developmental checkpoints guarded by regulated necrosis. Cell. Mol. Life Sci. 73, 2125–2136.

10. Dolle, P., Ruberte, E., Kastner, P., Petkovich, M., Stoner, C.M., Gudas, L.J., and Chambon, P. (1989). Differential expression of genes encoding alpha, beta and gamma retinoic acid receptors and CRABP in the developing limbs of the mouse. Nature 342, 702–705.

11. Duester, G. (2008). Retinoic acid synthesis and signaling during early organogenesis. Cell 134, 921–931.

12. Durand, B., Saunders, M., Leroy, P., Leid, M., and Chambon, P. (1992). All-trans and 9-cis retinoic acid induction of CRABPII transcription is mediated by RAR-RXR heterodimers bound to DR1 and DR2 repeated motifs. Cell 71, 73–85.

13. Green, D. (2005). Apoptotic pathways: ten minutes to dead. Cell 121, 671–674.

14. Harima, Y., Takashima, Y., Ueda, Y., Ohtsuka, T., and Kageyama, R. (2013). Accelerating the tempo of the segmentation clock by reducing the number of introns in the Hes7 gene. Cell Rep. 3, 1–7.

15. He, S., Wang, L., Miao, L., Wang, T., Du, F., Zhao, L., and Wang, X. (2009). Receptor interacting protein kinase-3 determines cellular necrotic response to TNF-alpha. Cell 137, 1100–1111.

16. Hitomi, J., Christofferson, D.E., Ng, A., Yao, J., Degterev, A., Xavier, R.J., and Yuan, J. (2008). Identification of a molecular signaling network that regulates a cellular necrotic cell death pathway. Cell 135, 1311–1323.

17. Horton, C., and Maden, M. (1995). Endogenous distribution of retinoids during normal development and teratogenesis in the mouse embryo. Dev. Dyn. 202, 312–323.

18. Itoh, N., Yonehara, S., Ishii, A., Yonehara, M., Mizushima, S., Sameshima, M., Hase, A., Seto, Y., and Nagata, S. (1991). The polypeptide encoded by the cDNA for human cell surface antigen Fas can mediate apoptosis. Cell, 66, 233–243.

19. Kaiser, W.J., Upton, J.W., Long, A.B., Livingston-Rosanoff, D., Daley-Bauer, L.P., Hakem, R., Caspary, T., and Mocarski, E.S. (2011). RIP3 mediates the embryonic lethality of caspase-8-deficient mice. Nature 471, 368–372.

20. Kikuchi, M., Kuroki, S., Kayama, M., Sakaguchi, S., Lee, K.K., and Yonehara, S. (2012). Protease activity of procaspase-8 is essential for cell survival by inhibiting both apoptotic and nonapoptotic cell death dependent on receptor-interacting protein kinase 1 (RIP1) and RIP3. J. Biol. Chem. 287, 41165–41173.

21. Kobayashi, Y., and Yonehara, S. (2009). Novel cell death by downregulation of eEF1A1 expression in tetraploids. Cell Death Differ. 16, 139–150.

22. le Maire, A., Teyssier, C., Erb, C., Grimaldi, M., Alvarez, S., de Lera, A.R., Balaguer, P., Gronemeyer, H., Royer, C.A., Germain, P., Bourguet, W. (2010). A unique secondary-structure switch controls constitutive gene repression by retinoic acid receptor. Nat. Struct. Mol. Biol. 17, 801–807.

23. Lee, S., Lee, B., Lee, J.W., and Lee, S.K. (2009). Retinoid signaling and neurogenin2 function are coupled for the specification of spinal motor neurons through a chromatin modifier CBP. Neuron 62, 641–654.

24. Lee, T.I., Johnstone, S.E., and Young, R.A. (2006). Chromatin immunoprecipitation and microarray-based analysis of protein location. Nat. Protoc. 1, 729–748.

25. Loudig, O., Babichuk, C., White, J., Abu-Abed, S., Mueller, C., and Petkovich, M. (2000). Cytochrome P450RAI(CYP26) promoter: a distinct composite retinoic acid response element underlies the complex regulation of retinoic acid metabolism. Mol. Endocrino. 14, 1483–1497.

26. Mandal, P., Berger, S.B., Pillay, S., Moriwaki, K., Huang, C., Guo, H., Lich, J.D., Finger, J., Kasparcova, V., Votta, B., et al. (2014). RIP3 induces apoptosis independent of pronecrotic kinase activity. Mol. Cell 56, 481–495.

27. Mangelsdorf, D.J., Ong, E.S., Dyck, J.A., and Evans, R.M. (1990). Nuclear receptor that identifies a novel retinoic acid response pathway. Nature, 345, 224–229.

28. Mangelsdorf, D.J., Umesono, K., Kliewer, S.A., Borgmeyer, U., Ong, E.S., and Evans, R.M. (1991). A direct repeat in the cellular retinol-binding protein type II gene confers differential regulation by RXR and RAR. Cell, 66, 555–561.

29. Mark, M., Ghyselinck, N.B., Chambon, P. (2005). Function of retinoid nuclear receptors: lessons from genetic and pharmacological dissections of the retinoic acid signaling pathway during mouse embryogenesis. Annu. Rev. Pharmacol. Toxicol. 46, 451–480.

30. Mattei, M.G., Riviere, M., Krust, A., Ingvarsson, S., Vennstrom, B., Islam, M.Q., Levan, G., Kautner, P., Zelent, A., Chambon, P., et al. (1991). Chromosomal assignment of retinoic acid receptor (RAR) genes in the human, mouse, and rat genomes. Genomics 10, 1061–1069.

31. Minamida, Y., Someda, M., and Yonehara, S. (2014). FLASH/casp8ap2 is indispensable for early embryogenesis but dispensable for proliferation and differentiation of ES cells. PLoS One 9, e108032.

32. Moutier, E., Ye, T., Choukrallah, M.A., Urban, S., Osz, J., Chatagnon, A., Delacroix, L., Langer, D., Rochel, N., Moras, D., et al. (2012) Retinoic acid receptors recognize the mouse genome through binding elements with diverse spacing and topology. J. Biol. Chem. 287, 26328–26341.

33. Muzio, M., Chinnaiyan, A. M., Kischkel, F. C., O’Rourke, K., Shevchenko, A., Ni, J., Scaffidi, C., Bretz, J. D., Zhang, M., Gentz, R., et al. (1996). FLICE, a novel FADD-homologous ICE/CED-3-like protease, is recruited to the CD95 (Fas/APO-1) death-inducing signaling complex. Cell 85, 817–827.

34. Oberst, A., Dillon, C.P., Weinlich, R., McCormick, L.L., Fitzgerald, P., Pop, C., Hakem, R., Salvesen, G.S., and Green, D.R. (2011). Catalytic activity of the caspase-8-FLIP(L) complex inhibits RIPK3-dependent necrosis. Nature 471, 363–367.

35. Ogura, T., and Evans, R.M. (1995). A retinoic acid-triggered cascade of HOXB1 gene activation. Proc. Natl. Acad. Sci. USA 92, 387–391.

36. Rhinn, M., and Dolle, P. (2012). Retinoic acid signalling during development. Development 139, 843–858.

37. Rossant, J., Zirngibl, R., Cado, D., Shago, M., and Giguere, V. (1991). Expression of a retinoic acid response element-hsplacZ transgene defines specific domains of transcriptional activity during mouse embryogenesis. Genes Dev. 5, 1333–1344.

38. Sakamaki, K., Inoue, T., Asano, M., Sudo, K., Kazama, H., Sakagami, J., Sakata, S., Ozaki, M., Nakamura, S., Toyokuni, S., et al. (2002). Ex vivo whole-embryo culture of caspase-8-deficient embryos normalize their aberrant phenotypes in the developing neural tube and heart. Cell Death Differ. 9, 1196–1206.

39. Salvesen, G. S., and Dixit, V. M. (1997). Caspases: intracellular signaling by proteolysis. Cell 91, 443–446.

40. Thornberry, N. A., and Lazebnik, Y. (1998). Caspases: enemies within. Science 281, 1312–1316.

41. Um, S., Harbers, M., Benecke, A., Pierrat, B., Losson, R., and Chambon, P. (1998). Retinoic acid receptors interact physically and functionally with the T:G mismatch-specific thymine-DNA glycosylase. J. Biol. Chem. 273, 20728–20736.

42. Varfolomeev, E.E., Schuchmann, M., Luria, V., Chiannilkulchai, N., Beckmann, J.S., Mett, I. L., Rebrikov, D., Brodianski, V.M., Kemper, O.C., Kollet, O., et al. (1998). Targeted disruption of the mouse Caspase 8 gene ablates cell death induction by the TNF receptors, Fas/Apo1, and DR3 and is lethal prenatally. Immunity 9, 267–276.

43. Wang, H., Sun, L., Su, L., Rizo, J., Liu, L., Wang, L.F., Wang, F.S., and Wang, X. (2014). Mixed lineage kinase domain-like protein MLKL causes necrotic membrane disruption upon phosphorylation by RIP3. Mol. Cell 54, 133–146.

44. Wang, Z.Y., and Chen, Z. (2008). Acute promyelocytic leukemia: From highly fatal to highly curable. Blood 111, 2505–2515.

45. Watanabe, K., Kamiya, D., Nishiyama, A., Katayama, T., Nozaki, S., Kawasaki, H., Watanabe, Y., Mizuseki, K., and Sasai, Y. (2005). Directed differentiation of telencephalic precursors from embryonic stem cells. Nat. Neurosci. 8, 288–296.

46. Yang, Y., Ma, J., Chen, Y., and Wu, M. (2004). Nucleocytoplasmic shuttling of receptor-interacting protein 3 (RIP3): identification of novel nuclear export and import signals in RIP3. J. Biol. Chem. 279, 38820–38829.

47. Yonehara, S., Ishii. A., and Yonehara, M. (1989). A cell-killing monoclonal antibody (anti-Fas) to a cell surface antigen co-downregulated with the receptor of tumor necrosis factor. J Exp Med, 169, 1747–1756.

48. Zelent, A., Krust, A., Petkovich, M., Kastner, P., and Chambon, P. (1989). Cloning of murine alpha and beta retinoic acid receptors and a novel receptor gamma predominantly expressed in skin. Nature 339, 714–717.

49. Zhang, D.W., Shao, J., Lin, J., Zhang, N., Lu, B.J., Lin, S.C., Dong, M.Q., and Han, J. (2009). RIP3, an energy metabolism regulator that switches TNF-induced cell death from apoptosis to necrosis. Science 325, 332–336.

